# Interactions of pathogenic and commensal strains of Mannheimia haemolytica with differentiated bovine airway epithelial cells grown at an air-liquid interface

**DOI:** 10.1101/197947

**Authors:** Daniel Cozens, Erin Sutherland, Miquel Lauder, Geraldine Taylor, Catherine C. Berry, Robert L. Davies

## Abstract

*Mannheimia haemolytica* serotype A2 is a common commensal species present in the nasopharynx of healthy cattle. However, prior to the onset of bovine pneumonic pasteurellosis, there is sudden increase in *M. haemolytica* serotype A1 within the upper respiratory tract. The events during this selective proliferation of serotype A1 strains are poorly characterised. In this investigation, a differentiated bovine airway epithelial cell culture was used to study the interactions of A1 and A2 bovine isolates with the respiratory epithelium. This model reproduced the key defences of the airway epithelium, including tight junctions and mucociliary clearance. Although initial adherence of the serotype A1 strains was low, by 12 hours post-infection the bacteria was able to traverse the tight junctions to form foci of infection below the apical surface. The size, density and number of these foci increased with time, as did the cytopathic effects observed in the bovine bronchial epithelial cells. Penetration of *M. haemolytica* A1 into the sub-apical epithelium was shown to be through transcytosis but not paracytosis. Commensal A2 bovine isolates however were not capable of colonising the model to a high degree, and did not penetrate the epithelium following initial adherence at the apical surface. This difference in their ability to colonise the respiratory epithelium may account for the sudden proliferation of serotype A1 in the onset of pneumonia pasteurellosis. The pathogenesis observed was replicated by virulent A2 ovine isolates; however colonisation was 10-fold lower in comparison to bovine A1 strains. This investigation provides new insight into the interactions of *M. haemolytica* with bovine airway epithelial cells which are occurring *in vivo* during pneumonia pasteurellosis.

## Introduction

Bovine respiratory disease (BRD) is a multifactorial condition of cattle that causes significant economic losses (>$3 billion annually in the USA alone) to the cattle industry worldwide [1-3]. The pathogenesis of BRD is complex, involving poorly understood interactions between various viral and bacterial pathogens and the host; environmental stress is also an important pre-disposing factor leading to the outbreak of disease [3-6]. Although many of the viral and bacterial pathogens can potentially cause disease themselves, it is generally accepted that viral infection often occurs first and predispose cattle to subsequent bacterial infection [3-9]. Pneumonic pasteurellosis is one of the most severe forms of BRD; it is characterized by an acute lobar fibronecrotizing pneumonia or pleuropneumonia and is associated with the bacterial pathogen *M. haemolytica* [3, 5, 10, 11].

*Mannheimia haemolytica* occurs naturally as a commensal in the upper respiratory tract of healthy cattle [12, 13] but, under circumstances described above, is frequently associated with disease [1, 10, 12]. The bacterium comprises 12 capsular serotypes [14] but it is widely recognized that serotype A2 strains are most commonly associated with healthy cattle, where they reside as commensals in the nasopharynx and tonsils; conversely, serotype A1 (and more recently A6) strains are mainly responsible for disease [1, 3, 10, 15, 16]. However, in addition to differences in capsular polysaccharide biochemistry and structure [17, 18], serotype A1/A6 and A2 strains of *M. haemolytica* represent distinct chromosomal genotypes [19, 20] and can also be distinguished by differences in their outer membrane protein (OMP) profiles [21], lipopolysaccharide types [21, 22] and nucleotide sequence variation in various virulence-associated genes including *lktA* [23], *ompA* [24], *tbpA* and *tbpB* [25], *plpE* [26] as well as a number of other genes [20]. The upper respiratory tract of healthy cattle is predominantly colonized by serotype A2 strains but, for reasons that are not clear (but probably related to stress and/or viral infection), a transition occurs within this microenvironment which leads to a sudden explosive proliferation in the number of serotype A1/A6 bacteria present and subsequent colonization [1, 3]. This sudden and selective explosion in the A1/A6 population within the upper respiratory tract leads to the inhalation of bacteria-containing aerosol droplets into the trachea and lungs and the onset of pneumonic pasteurellosis [27]. Crucially, the specific bacterial and host factors responsible for the sudden shift from commensal serotype A2 to pathogenic serotype A1/A6 populations within the upper respiratory tract are not clear.

The leukotoxin (LktA) of *M. haemolytica* plays a central role in the pathogenesis of pneumonic pasteurellosis and significant attention has been given to understanding the molecular mechanisms associated with LktA activity within the lung [1, 3, 28, 29]. In contrast, there has been far less focus on the interactions of *M. haemolytica* with respiratory airway epithelial cells and events that might account for the sudden proliferation of serotype A1/A6 bacteria within the upper respiratory tract. A contributing factor to our poor understanding of early host-pathogen interactions associated with pneumonic pasteurellosis, and indeed BRD in general, is the lack of physiologically-relevant and reproducible methodologies with which to study the intricate molecular and immunological interactions between pathogens and host. Traditionally, submerged, two-dimensional cultures of a single cell type have been used to investigate interactions of *M. haemolytic* and other BRD pathogens within the bovine respiratory tract [30-32] but these have numerous limitations: they do not reflect the multicellular complexity of the parental tissue *in vivo*, they lack its three-dimensional (3-D) architecture, and the physiological conditions are not representative of those found within the respiratory tract. However, these characteristics that are lacking in submerged cultures can be recapitulated using differentiated airway epithelial cells (AECs) grown at an air-liquid interface (ALI) and, in recent years, such cell culture approaches have been used to study the interactions of various bacterial and viral pathogens with different host species [33-43].

We have previously investigated the growth conditions required for optimal growth and differentiation of bovine bronchial epithelial cells at an ALI [44] and assessed the temporal differentiation of these cells to identify an optimum window suitable for infection studies [45]. The aim of the present study was to investigate the interactions of a panel of *M. haemolytica* isolates, representing virulent and commensal strains recovered from both cattle and sheep, with differentiated bovine bronchial epithelial cells grown at an ALI. The course of infection was followed for up to five days using various microscopic approaches and the production of selected cytokines measured to ascertain the epithelial cell response.

## Materials and Methods

### Bacterial cultures

Eight wild-type *M. haemolytica* strains (Table 1) isolated from both cattle and sheep were included in this investigation. The strains were isolated from either pneumonic or healthy animals. Bacteria were routinely grown on brain-heart infusion (BHI) agar supplemented with 5% (v/v) defibrinated sheep blood overnight at 37 °C. Broth cultures were grown in BHI broth at 37 °C with agitation.

**Table 1.**
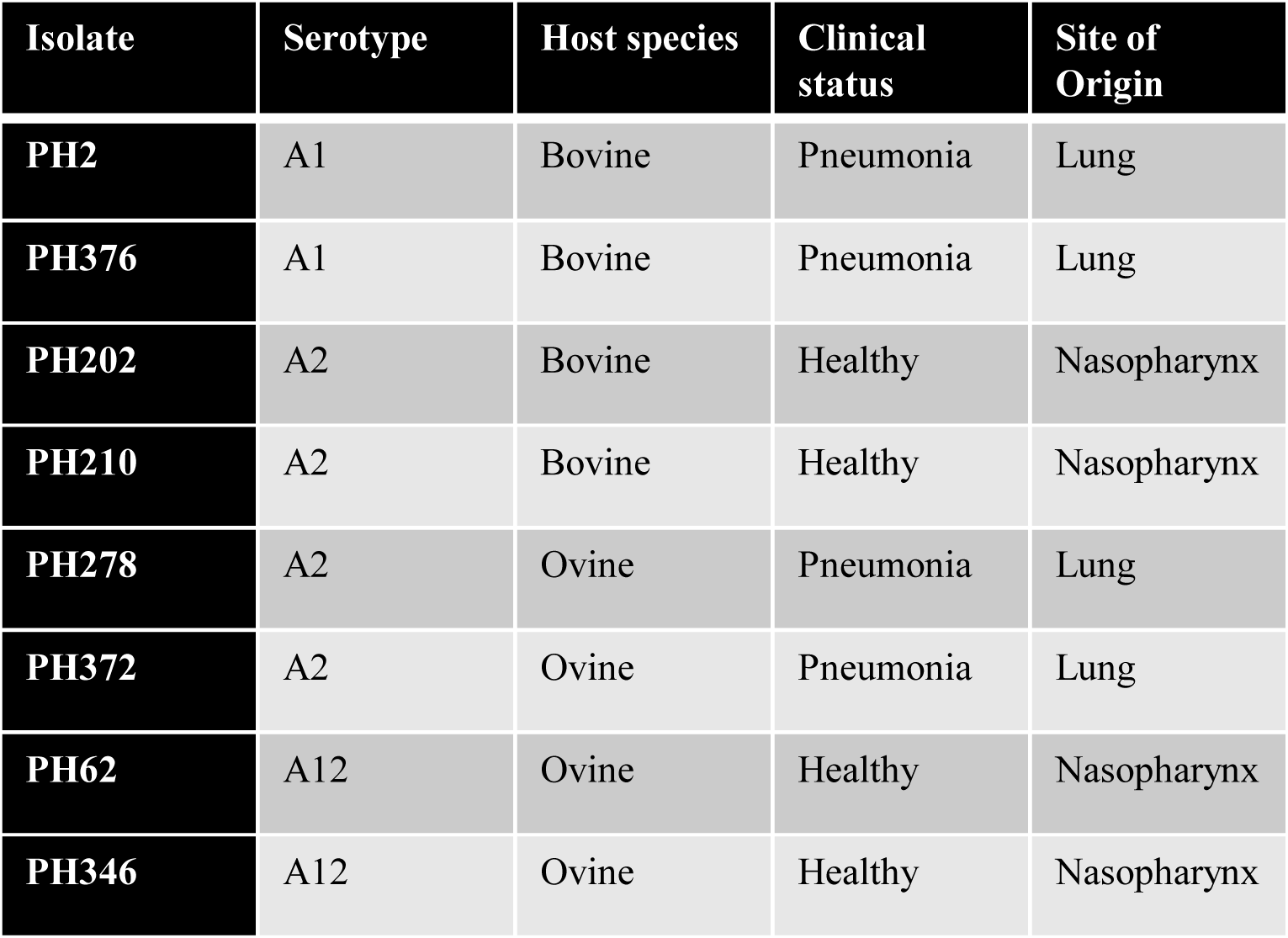
*M. haemolytica* strains utilised in this investigation.

### Culture of bovine bronchial epithelial cells

Bronchial epithelial cells were isolated from cattle aged 24-30 months, as described previously [44]. Tissue was collected from cattle immediately post-slaughter at Sandyford Abattoir Ltd., UK. The bronchial tract was swabbed to ensure there was no pre-existing bacterial or fungal infection. *Ex vivo* bronchi tissue was also collected, fixed in 2% (w/v) formaldehyde and sectioned for histological analysis to confirm the health of the donor animal. Briefly, the main and lobar bronchi were dissected from the lungs and the surrounding tissue removed. The BBECs were isolated from the epithelium by incubation overnight at 4°C in ‘digestion medium’ composed of Dulbecco’s modified Eagle’s medium (DMEM) and Ham’s nutrient F-12 (1:1) containing 1 mg/ml dithioreitol, 10 µg/ml DNAase and 1 mg/ml Protease XIV from *Streptomyces griseus*, supplemented with penicillin (100 U/ml), streptomycin (100 µg/ml) and amphotericin (2.5 µg/ml) (Sigma-Aldrich). All subsequent media, with the exception of media utilised during infection of the BBEC cultures were also supplemented with penicillin-streptomycin and amphotericin. Digestion of the bronchial epithelium was halted by the addition of foetal calf serum to give a final concentration of 10% (v/v). Rigorous rinsing of the luminal surface was used to remove loosely-attached epithelial cells. The resulting suspension was centrifuged and resuspended in ‘submerged growth medium’ (SGM), comprised of DMEM/Ham’s F-12 (1:1) supplemented with 10% (v/v) foetal calf serum. Cells were seeded into T75 tissue culture flasks (5 x 10^6^ cells/flask) for expansion. The flasks were incubated at 37°C in 5% CO_2_ and 14% O_2_, in a humidified atmosphere. At 80-90 % confluency (~4 days post-seeding) the flasks of BBECS were harvested. Cells were detached using 0.25% trypsin-EDTA solution, centrifuged and resuspended in SGM at a density of 5 × 10^5^ cells/ml. The BBECS were seeded into the apical chamber of tissue culture inserts (Thincerts, Greiner #66540, polyethylene terephthalate membrane, 0.4 µm pore diameter, 1 × 10^8^ pore per cm^2^) at a density of 2.5 × 10^5^ cells per insert. Cultures were incubated at 37 °C, 5% CO_2_, 14% O_2_, in a humidified atmosphere. Following overnight incubation, the apical medium of the culture was removed and the apical surface washed with 0.5 ml PBS to remove unattached cells. The SGM media in the apical and basolateral compartments was then replaced. This process was repeated every 2 – 3 days. The trans-epithelial electrical resistance (TEER) of the cultures were monitored on a daily basis using an EVOM2 epithelial voltohmmeter (World Precision Instruments, UK), as per the manufacturer’s instruction. Once the TEER reached above 200 Ω/cm^2^ (~2 days post-seeding) the SGM was replaced with a mixture of SGM and ‘air-liquid interface medium’ (ALIM) (1:1). The ALIM was composed of DMEM and airway epithelial cell growth medium (Promocell) (1:1) supplemented with 10 ng/ml epidermal growth factor, 100 nM retinoic acid, 6.7 ng/ml triiodothyronine, 5 µg/ml insulin, 4 µl/ml bovine pituitary extract, 0.5 µg/ml hydrocortisone, 0.5 µg/ml epinephrine and 10 µg/ml transferrin (all Promocell). When the TEER value was above 500 Ω cm^2^ (~6 days post-seeding), an ALI was generated by removing the medium in the apical compartment, thereby exposing the epithelial cells to the atmosphere (day 0 post-ALI). Following the formation of the ALI, the cells were fed exclusively from the basal compartment with ALIM. Apical washing, basal feeding and TEER measurements were performed every 2 - 3 days until day 21 post-ALI.

### Infection of bovine bronchial epithelial cells

The BBEC cultures were infected on day 21 post-ALI. The apical and basal compartments were washed twice with PBS, 24 hours prior to infection. The cultures were subsequently fed with 1 ml ALIM with the omission of penicillin-streptomycin and amphotericin. Bacteria used in the infection were collected from fresh overnight plate cultures, grown in BHI broth to exponential phase, and resuspended in PBS at 10^9^ cfu/ml. The bacterial suspension was used to inoculate the BBEC cultures apically. Each insert was inoculated with 25 µl of bacterial suspension (2.5 x 10^7^ cfu/insert). Cultures were incubated at 37 °C until the stated time point post-infection. For infection of undifferentiated BBECs, cultures were infected at day 0 post-ALI.

### Quantification of bacterial adhesion

At stated time points following infection, a viability count was performed on the adherent *M. haemolytica*. The ALIM was removed from the basal compartment and the apical surface of the transwell was washed three times with 1 ml PBS. The three washes were subsequently pooled and the number of viable bacteria was also assessed. The BBEC were incubated in 0.5 ml PBS with 1% Triton X-100 to permeablise the epithelial cells. The membrane was scraped to mechanical disintegrate the culture. Viable bacteria in the lysate and apical washes were quantified using 10-fold serial dilutions, performed in triplicate, and plating on BHI agar with 5% (v/v) defibrinated sheep blood, using the Miles and Misra method. Plates were incubated for six hours at 37 °C and the number of colony-forming units (CFU) counted. Bacterial number was expressed as a percentage of the inoculum. For the gentamicin protection assay, the apical surface was treated with 200 µg/ml gentamicin for one hour at 37 °C prior to permabilisation. For each animal, bacterial adherence was quantified in three independent BBEC cultures at all time points.

### Histology and immunohistochemistry

At the stated time points post-infection, cultures were fixed by incubation with 4% (w/v) paraformaldehyde for 15 min at room temperature and rinsed in PBS. The samples were subsequently dehydrated using a series of increasing ethanol concentrations, cleared with xylene and infiltrated with paraffin wax. Sections of the wax blocks were cut at 2.5 µm thickness using a Thermoshandon Finesse ME+ microtome. Samples were stained with haematoxylin and eosin (H&E) using standard histological techniques. Further sections were stained for immunohistochemistry. Rabbit anti-bovine OmpA antibody was used to identify bovine *M. haemolytica* strains and rabbit anti-ovine OmpA antibody was used to identify ovine *M. haemolytica* strains, at a dilution of 1:800. Heat-induced epitope retrieval was performed using a Menarini Access Retrieval Unit and staining conducted using a Dako Autostainer. Endogenous peroxidase was blocked with 0.3% (v/v) H_2_O_2_ in PBS. Following incubation with the primary antibody, binding was identified by application of an anti-rabbit HRP-labelled polymer and visualization with a REAL EnVision Peroxidase/DAB+ Detection System (Dako; #K3468). Samples were subsequently counterstained with Gill’s haematoxylin, dehydrated, cleared and mounted in synthetic resin before sectioning. Tissue sections were viewed with a Leica DM2000 microscope.

### Immunofluorescence microscopy

At the stated time points post-infection, cultures were fixed by incubation with 4% (w/v) paraformaldehyde for 15 min at room temperature and rinsed in PBS. Samples were immunofluorescently stained as previously described[44]. Briefly, samples were permeablised using permabilization buffer (PBS with 0.5% [v/v] Triton X-100, 100 ml/ml sucrose, 4.8 mg/ml HEPES, 2.9 mg/ml NaCl and 600 μg/ml MgCl_2_, pH 7.2) for 10 min at room temperature. Samples were blocked by incubation with PBS containing 0.05% (v/v) Tween-20, 10% (v/v) goat serum and 1% (w/v) bovine serum albumin. The primary-secondary antibody pairings were applied as follows. Bovine *M. haemolytica* strains were detected using *M. haemolytica* antisera produced in cattle (1:50 dilution) and visualised with goat anti-bovine-FITC antibody (1:400, Thermo Fisher #A18752). Ovine *M. haemolytica* strains were detected using rabbit anti-ovine OmpA antibody (1:50 dilution) and visualised with goat anti-rabbit-Alexa Fluor 488 (1:400 dilution; Thermo Fisher; #A-11008). Ciliated cells were detected with mouse anti-β-tubulin antibody (1:50 dilution; Abcam; #ab131205). Tight-junction formation was detected with mouse anti-ZO-1 antibody (1:50 dilution; Thermo Fisher; #33910). Both anti-β-tubulin and anti-ZO-1 antibody binding was detected with anti-mouse-Alexa Fluor 568 (1:400 dilution; Thermo Fisher; #A-11031). The cultures were incubated with antibodies diluted in blocking buffer for 1 h at room temperature. Samples were washed three times in PBS containing 0.05% (v/v) Tween-20 for 2 min following each incubation. Blocking was repeated after each primary-secondary pairing. Nuclei were stained with 300 nM 4',6 diamidino-2-phenylindole (DAPI) for 10 min. Following staining, membranes were cut from their insert and mounted in Vectashield mounting medium (Vector Laboratories). Samples were observed on a Leica DMi8 microscope. Z-stack orthological representation was observed on a Zeiss AxioObserver Z1spinning disk confocal microscope. Analysis of captured images was performed using ImageJ software.

### Scanning electron microscopy

At the stated time points following infection, cultures were fixed in 1.5% (v/v) glutaraldehyde in 0.1 M sodium cacodylate buffer for 1 h at 4°C. Samples were subsequently rinsed three times with 0.1 M sodium cacodylate buffer and post-fixed in 1% (w/v) osmium tetroxide for 1 h at room temperature. The cultures were washed three times for 10 min with distilled water, stained with 0.5% (w/v) uranyl acetate for 1 h in the dark, washed twice with distilled water and dehydrated through a series of increasing ethanol concentrations. The samples were further dehydrated in hexamethyldisilazane before being placed in a desiccator overnight. Membranes were cut from the inserts, mounted onto aluminium SEM stubs and gold sputter-coated. The cultures were analysed on a Jeol 6400 scanning electron microscope at 10 kV.

## Results

### *M. haemolytica* infection of undifferentiated bovine bronchial epithelial cells

The ability of *M. haemolytica* to adhere and colonise BBECs was first determined using undifferentiated cultures. These cultures consisted of primary isolated BBEC cultures grown in tissue culture inserts under submerged conditions. We have previously shown that under these conditions BBEC form undifferentiated monolayers. Staining for β-tubulin was indicative of cytoskeletal microtubules as opposed to cilial staining (Fig S1). However the cultures did possess tight junctions, as identified using marker Zona Occludens-1 (ZO-1) (Fig S2). The undifferentiated BBEC cultures were apically infected with either *M. haemolytica* strain PH2, an A1 serotype isolated from the lung of a pneumonic animal, or PH202, an A2 serotype isolated from the nasopharynx of a healthy animal. Adhesion and colonisation of the bacteria was quantified following infection (Fig 1A). Initial adherence at 0.5-2 hours post-infection (hpi) was comparable between the virulent PH2 strain and the commensal PH202 strain. Approximately 1% of the inoculum initially adhered to the BBECs (Fig 1A [i]). The majority of the inoculum was present in the apical washes (Fig 1A [ii]). At 24 hpi, there is a significant increase in the number of PH2 present in the culture, particularly the number of adherent bacteria (Two-way ANOVA), indicating that the strain was capable of highly colonising undifferentiated BBECs. This colonisation was not replicated by PH202; there was not a significant increase in either the number of adherent bacteria or bacteria removed from the monolayer in the apical wash.

**Figure 1.**
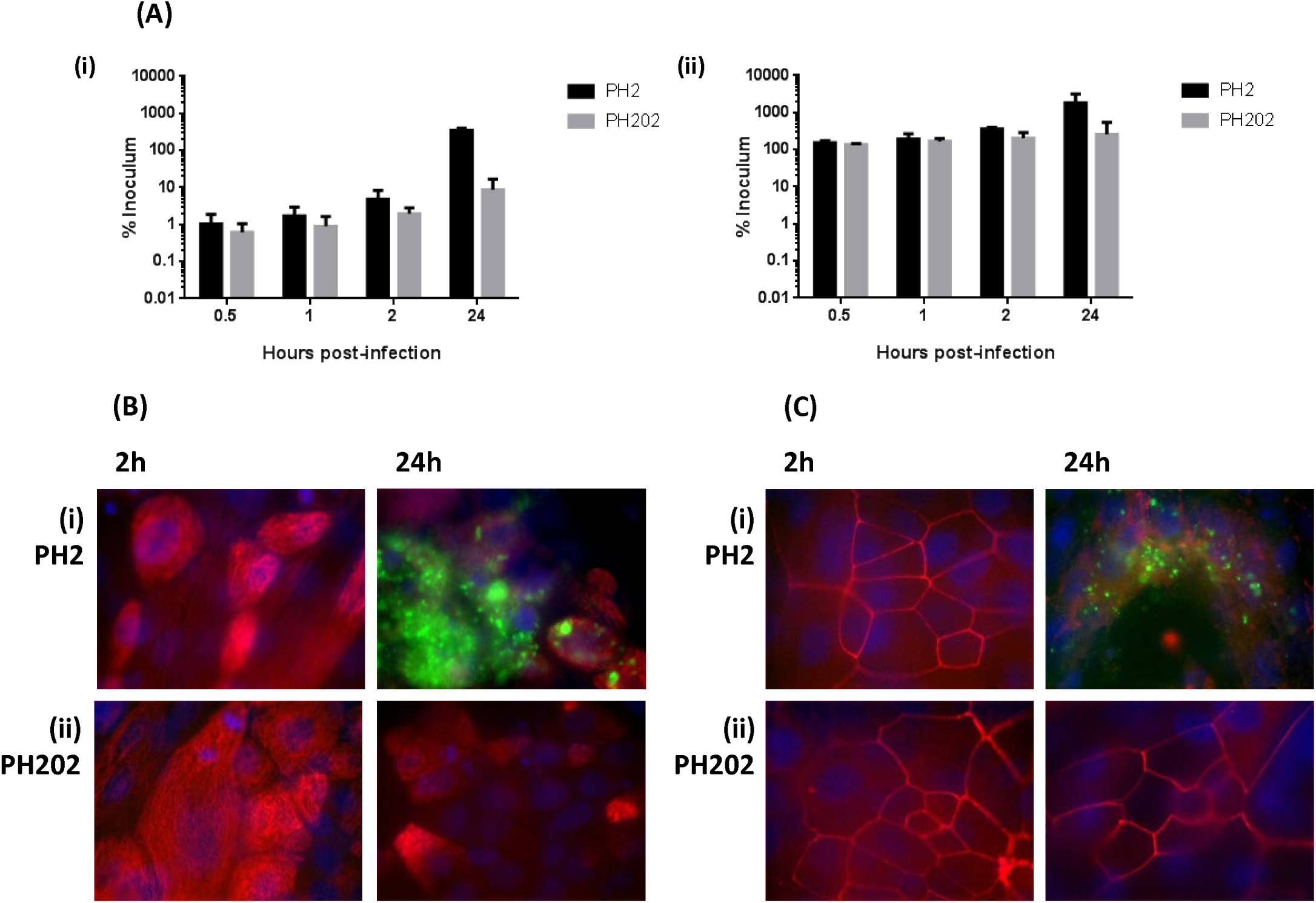
Infection of undifferentiated BBEC cultures by *M. haemolytica* strains. BBEC cultures were infected apically with *M. haemolytica* strains PH2 or PH202 (2.5 × 10^7^ cfu/insert) at day 0 post-ALI. At 0.5, 1, 2 and 24 hpi, cultures were apically washed to remove unbound bacteria, and colonisation assessed. (A) Quantification of the number of (i) adherent *M. haemolytica* and (ii) *M. haemolytica* present in the apical wash, as expressed as a percentage of the original inoculum. Three inserts were analysed per time point, and the data represents the mean +/- standard deviation from cultures derived from three different animals. (B-C) Cultures were fixed at the stated time post-infection and immunostained to detect colonisation of *M. haemolytica* with (B) β-tubulin (*M. haemolytica* - green; β-tubulin - red; nuclei – blue; x1000 magnification) or (C) tight junctions (*M. haemolytica* - green; ZO-1 -red; nuclei – blue; x1000 magnification). Representative images are shown of *M. haemolytica* strains (i) PH2 or (ii) PH202 at 2 and 24 hpi (see Fig S1 and S2).

The localisation of bacteria adherent to undifferentiated BBECs was detected by labelling with antisera raised against *M. haemolytica*. Epithelial cells were identified using β-tubulin and DAPI (Fig 1B and Fig S1). Between 0.5-2 hpi, adherence was low for both strains. A small population of bacteria was distributed on the apical surface of cells; however several cultures were visually devoid of adherent bacteria. Conversely, after 24 hpi, PH2 was near-confluent at numerous foci of infection present across the culture. These foci were separated by areas of much lower colonisation density. This pattern was not replicated by PH202, which at 24 hpi continued to show little to no evidence of adherence. Infected BBEC cultures were also labelled for tight junctional protein ZO-1 to identify the effect of *M. haemolytica* on tight junction integrity (Fig 1C). Tight junctions were shown to be stable across all time points following infection with PH2 and PH202. However, at the foci of infection, where PH2 was present at high number, tight junctions could not be observed. This was determined to be due to damage to the colonised epithelial cells resulting in a disruption in the integrity of the monolayer as opposed to direct targeting of the tight junction by PH2. This will be discussed in greater detail below.

### *M. haemolytica* infection of differentiated bovine bronchial epithelial cells

Adherence and colonisation of *M. haemolytica* was further determined using differentiated BBEC cultures. Primary BBEC were grown at an ALI in order to stimulate polarisation of airway epithelial cells into a culture which closely replicates the *in vivo* epithelium of the bovine respiratory tract (Fig 2). At 21 days post-ALI, the BBEC cultures were shown to form a pseudostratified columnar epithelium highly reminiscent to *ex vivo* tissue section (Fig 2A [i] & 2B [i]). The apical surface of the BBEC cultures displayed both a high degree of ciliation (Fig 2B [ii] & 2C [i]), and the formation of tight junction (Fig 2C [ii]), characteristic of the bovine airway lumen. This model has previously been well characterised and was shown to replicate other hallmarks of the airway epithelium, including the differentiation of mucus-producing goblet cells and active mucociliary clearance.

**Figure 2.**
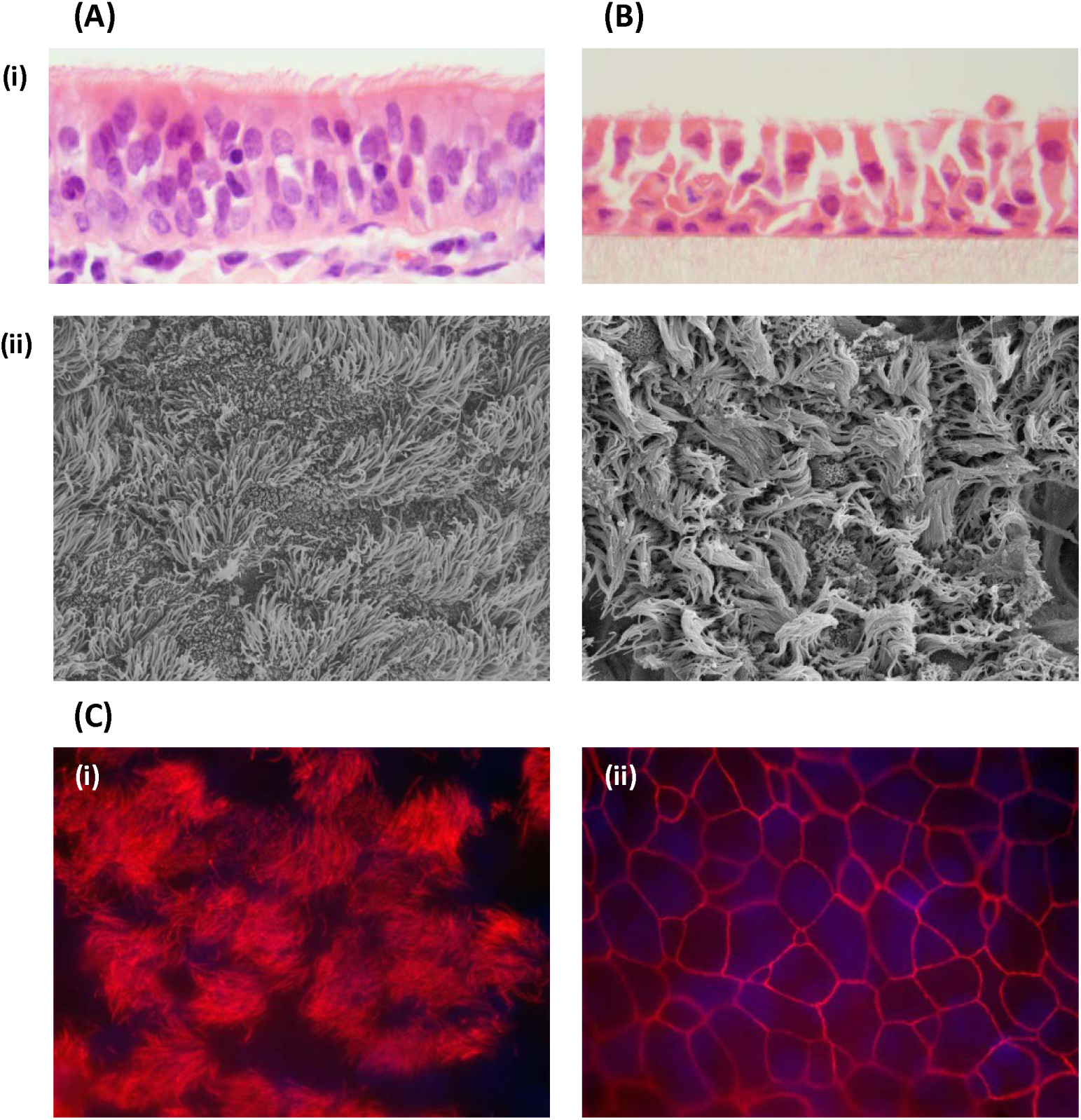
Differentiated BBEC cultures replicate the bovine bronchial epithelium. BBEC cultures were grown for 21 days at an ALI before fixation; sample of *ex vivo* tissue were also taken from the donor animal. Morphology was subsequently assessed using (A) examination by SEM (x2500 magnification) and (B) H&E stained histological sections (x1000 magnification). Representative images are shown of (i) *ex vivo* bovine bronchial epithelium and (ii) uninfected differentiated BBECs 21 days post-ALI. (C) BBEC cultures 21 days post-ALI were immunostained for markers of differentiation. Representative images are shown displaying (i) cilia formation (β-tubulin - red; nuclei – blue; x1000 magnification) and (ii) tight junction formation (ZO-1 - red; nuclei – blue; x1000 magnification).

The differentiated BBEC cultures were infected apically with either PH2 or PH202. The number of bacteria present in the culture was quantified using the Miles and Misra method at time points over a five day period (Fig 3). For both PH2 and PH202, initial adherence (0.5-2 hpi) was approximately 1% of the inoculum (Fig 3A); with the majority of the bacteria removed following apical washing of the BBEC culture (Fig 3B). The number of adherent PH2 increased within the BBEC cultures in a time-dependent manner from 6 hpi onwards (Fig 3A). At 24 hpi, the number of PH2 adherent to the model was approximately 1800% the initial inoculum. There was a subsequent decrease in the number of adherent PH2 at 48 hpi. It is hypothesised that this decrease was due to removal of damaged BBECs following apical washing, as discussed below. This increase in the number of adherent PH2 over time was accompanied by an increase in the number of bacteria removed in the apical wash (Fig 3B). Conversely, PH202 was not capable of colonising the BBEC model. In cultures derived from two of the three donor animals, no viable bacteria could be detected at 120 hpi. In BBEC cultures derived from a third animal, PH202 adhered to the culture at approximately 15% the initial inoculum, 100-fold lower than the virulent PH2 strain (Fig 3A). This trend is replicated in the number of bacteria removed by the apical wash (Fig 3B).

**Figure 3.**
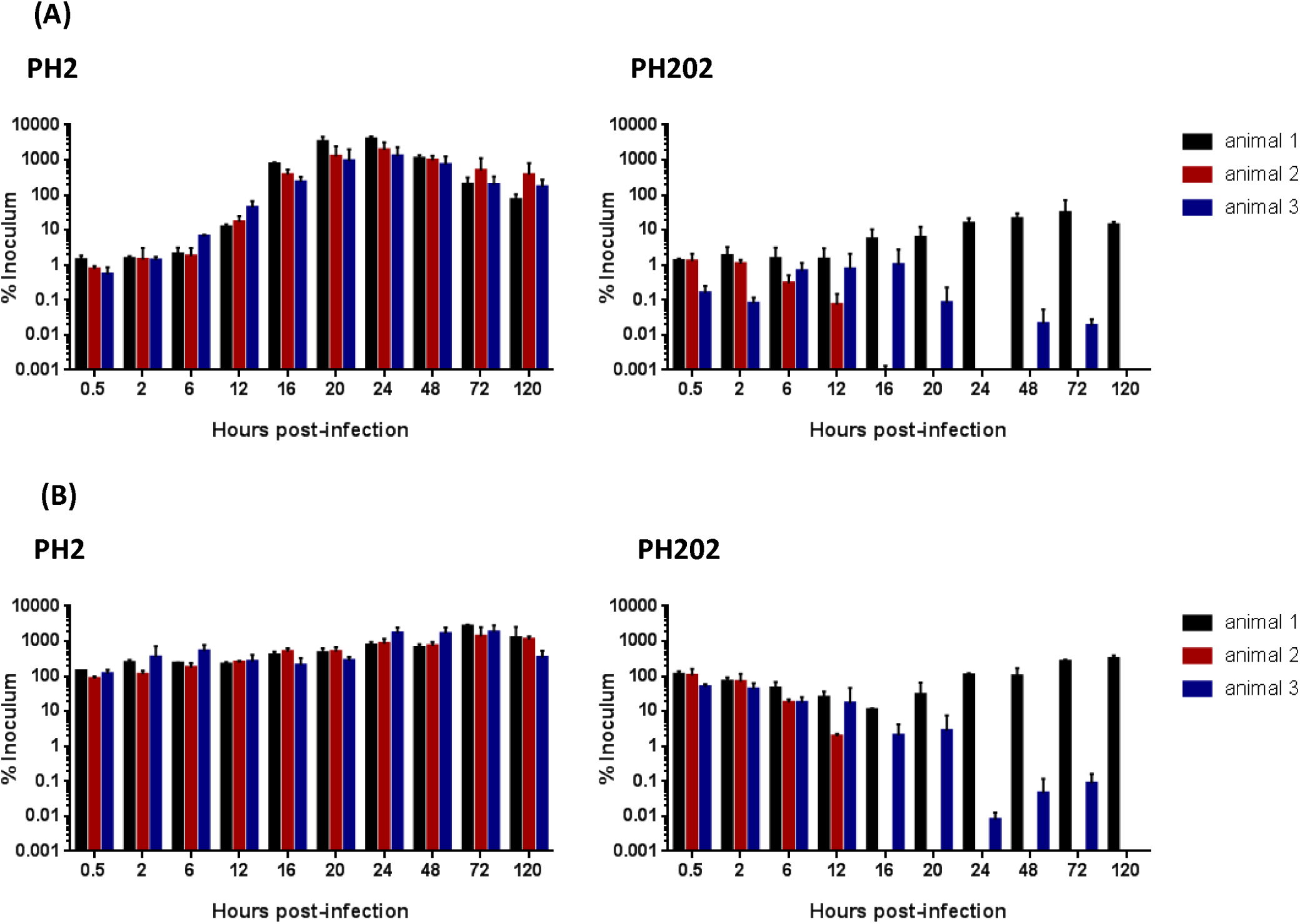
Adhesion of PH2 and PH202 to differentiated BBEC cultures. BBEC cultures were infected apically with *M. haemolytica* strains PH2 or PH202 (2.5 × 10^7^ cfu/insert) at day 21 post-ALI. At stated time points post-infection, cultures were apically washed to remove unbound bacteria. Quantification of the number of (A) adherent *M. haemolytica* and (B) *M. haemolytica* present in the apical wash per insert, as expressed as a percentage of the original inoculum. Three inserts were analysed per time point, and the data represents the mean +/-standard deviation.

### *M. haemolytica* form foci of infection in differentiated bovine bronchial epithelial cells

The distribution of *M. haemolytica* following infection of differentiated BBEC was determined using several microscopic techniques (Fig 4 & 5). The adhesion of bacteria to the apical surface was detected by labelling with antisera raised against *M. haemolytica*. From 0.5-6 hpi, PH2 was shown to be distributed across the apical surface of the culture at a low density (Fig 4A & S3). The bacteria did not display a preference for ciliated or non-ciliated cells during initial adherence. This observation was confirmed using SEM (Fig 4B & S4). At 12 hpi, PH2 became increasing abundant at the apical surface, forming focal areas of infection (Fig 4A [ii]). By this time point, PH2 had penetrated below the apical surface of the BBEC cultures, as shown in histological sections labelled for *M. haemolytica* using an anti-OmpA antibody (Fig 4C & S5). The density of PH2 at the foci of infection increased at 16 hpi (Fig 4A [iii]). These foci could be observed using SEM, in which PH2 was present in near-confluent consolidations below the apical surface (Fig 4B [iii]). Bacteria could not be observed at the apical surface in the proximity of the foci. Within the histological sections, PH2 at the foci were shown to have penetrated the entirety of the epithelium to the basal surface (Fig 4C [iii]). As exposure time increased, the number, size and density of the foci increased. This phenomenon coincides with an increased quantity of bacteria recovered from the apical surface (Fig 3A). By 24 hpi, foci were present at high numbers across the culture, with large regions heavily colonised by PH2 below the apical surface (Fig 4C [v]). The BBECs neighbouring the foci of infection did not display evidence of damage or cell death in the initial 24 hours following challenge by PH2.

**Figure 4.**
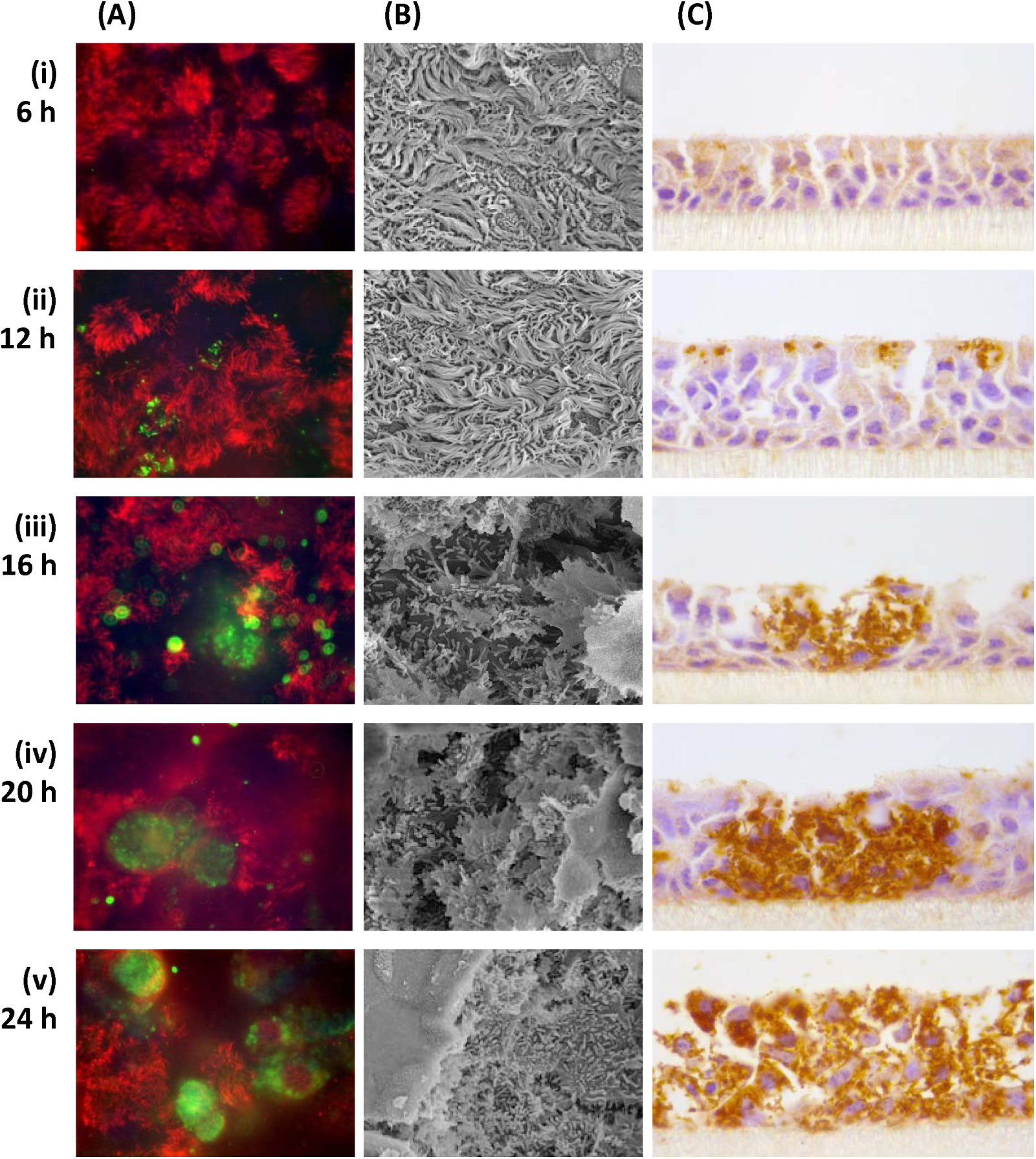
Infection of differentiated BBEC cultures by *M. haemolytica* PH2. BBEC cultures were infected apically with *M. haemolytica* strain PH2 (2.5 × 10^7^ cfu/insert) at day 21 post-ALI. At stated time points post-infection, cultures were apically washed to remove unbound bacteria, and fixed. Colonisation of PH2 was subsequently assessed using (A) immunofluorescence labelling of PH2 and cilia (*M. haemolytica* - green; β-tubulin - red; nuclei – blue; x1000 magnification), (B) examination by SEM (x2500 magnification) and (C) immunohistochemical-labelling of PH2 in histological sections (OmpA-labelled *M. haemolytica* stained brown; x1000 magnification). Representative images are shown of BBEC cultures at (i) 6, (ii) 12, (iii) 16, (iv) 20 and (v) 24 hpi (see Fig S3, S4 and S5).

**Figure 5.**
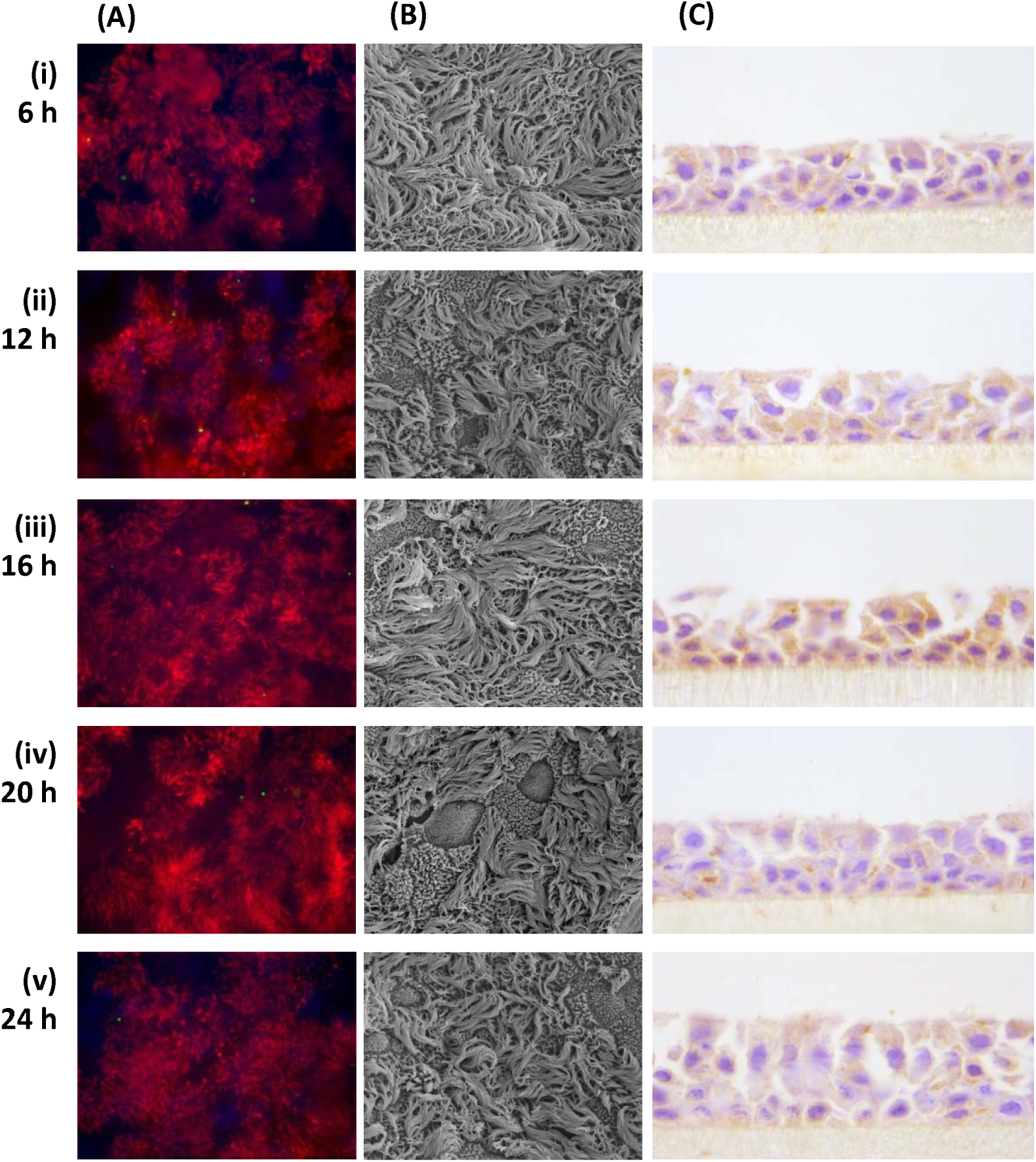
Infection of differentiated BBEC cultures by *M. haemolytica* PH202. BBEC cultures were infected apically with *M. haemolytica* strain PH202 (2.5 × 10^7^ cfu/insert) at day 21 post-ALI. At stated time points post-infection, cultures were apically washed to remove unbound bacteria, and fixed. Colonisation of PH202 was subsequently assessed using (A) immunofluorescence labelling of PH202 and cilia (*M. haemolytica* - green; β-tubulin - red; nuclei – blue; x1000 magnification), (B) examination by SEM (x2500 magnification) and (C) immunohistochemical-labelling of PH202 in histological sections (OmpA-labelled *M. haemolytica* stained brown; x1000 magnification). Representative images are shown of BBEC cultures at (i) 6, (ii) 12, (iii) 16, (iv) 20 and (v) 24 hpi (see Fig S3, S4 and S5).

This pattern of infection observed following challenge of differentiated BBEC cultures by virulent strain PH2 was not mimicked by commensal strain PH202 (Fig 5). The adherence of PH202 could often be barely detected on the apical surface using immunofluorescence labelling (Fig 5A) or SEM (Fig 5B). When PH202 was detected within the culture, the bacteria were present on the apical surface at a low population density. Histological sections of infected tissue confirmed that PH202 was not capable of penetrating the apical surface of the culture (Fig 5C).

The effect of apical infecting with PH2 or PH202 on the integrity of the BBEC cultures was investigated in histological sections (Fig 6 and S6). A semi-quantitative assessment of the degree of infection was conducted at each time point from cultures derived from three individual animals (Table 2). Evidence of infection following challenge by PH2 could be observed by 12 hpi, at which individual foci of infection could be observed in regions across the culture. The foci were increasingly abundant by 24 hpi. The BBECs at these foci displayed cytopathic effects. Airway epithelial cells colonised by high numbers of PH2 became increasingly rounded and detached from the epithelium (Fig 6 [v]). Infected cells displayed cytoskeletal staining for β-tubulin as opposed to cilial, indicating the cells are becoming dedifferentiated. By 48 hpi, the integrity of the epithelium was drastically reduced, and the majority of the culture was dislodged following apical washing (Fig S6). This may account for the reduction in CFU at 48 hpi (Fig 3A). Epithelial cells still attached to the epithelium appeared rounded (Fig S3) and were heavily colonised by bacteria (Fig S4). This observation was not replicated following infection with PH202. Cultures remained healthy until the time course was halted at 120 hpi (Fig S6). There was no evidence of cell rounding or increased cell death, with the exception of animal 1, which at 120 hpi showed reduced integrity of the cell layer. This was not observed in animal 2 or 3 (Table 2).

**Figure 6.**
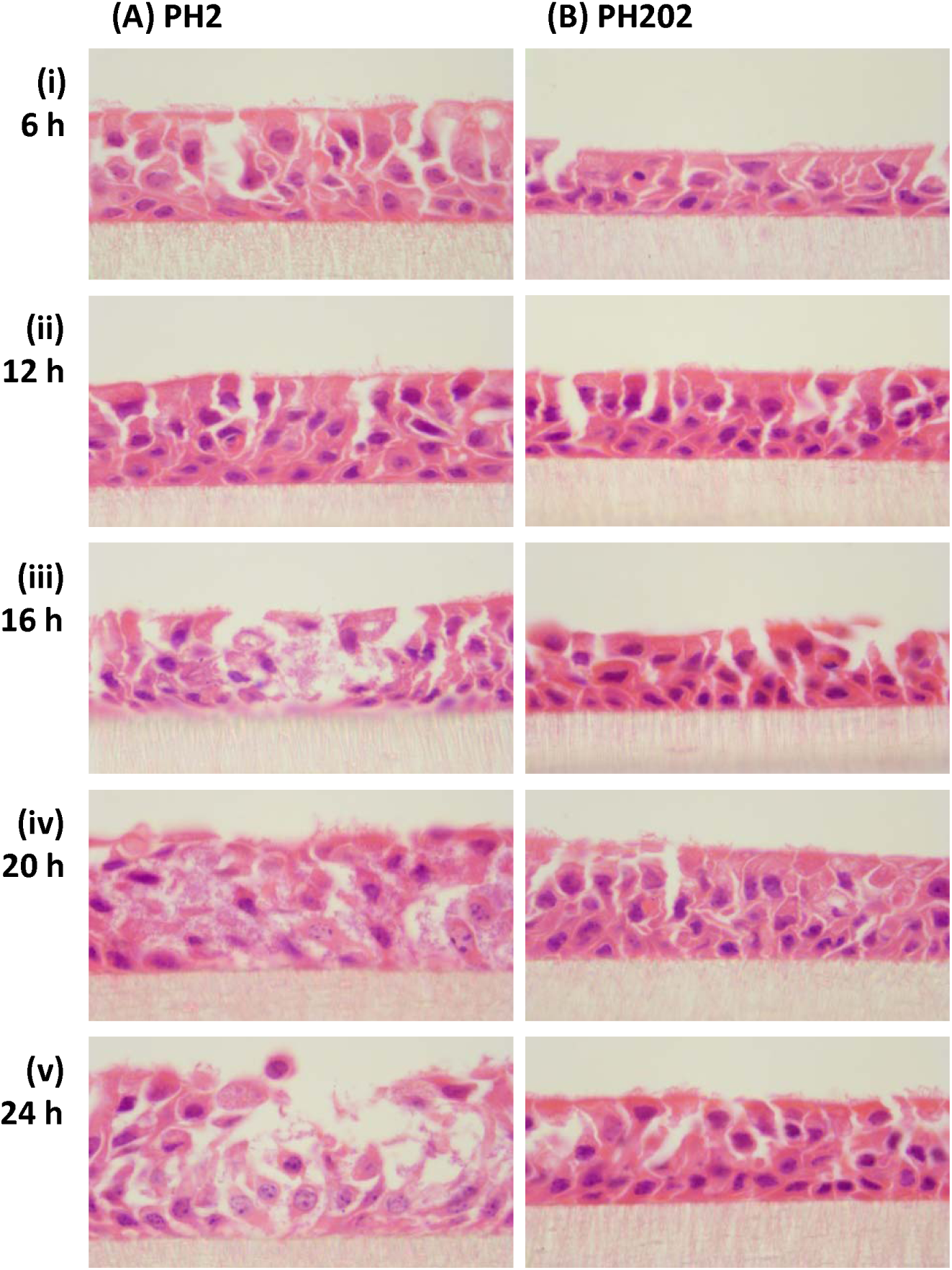
Histological assessment of *M. haemolytica* infection of differentiated BBEC cultures. BBEC cultures were infected apically with *M. haemolytica* strains (A) PH2 or (B) PH202 (2.5 × 10^7^ cfu/insert) at day 21 post-ALI. At stated time points post-infection, cultures were apically washed to remove unbound bacteria, fixed and paraffin-embedded using standard histological techniques. Sections were subsequently cut, deparaffinised and H&E stained. Representative images are shown of BBEC cultures at (i) 6, (ii) 12, (iii) 16, (iv) 20 and (v) 24 hpi (x1000 magnification; see Fig S6).

**Table 2.**
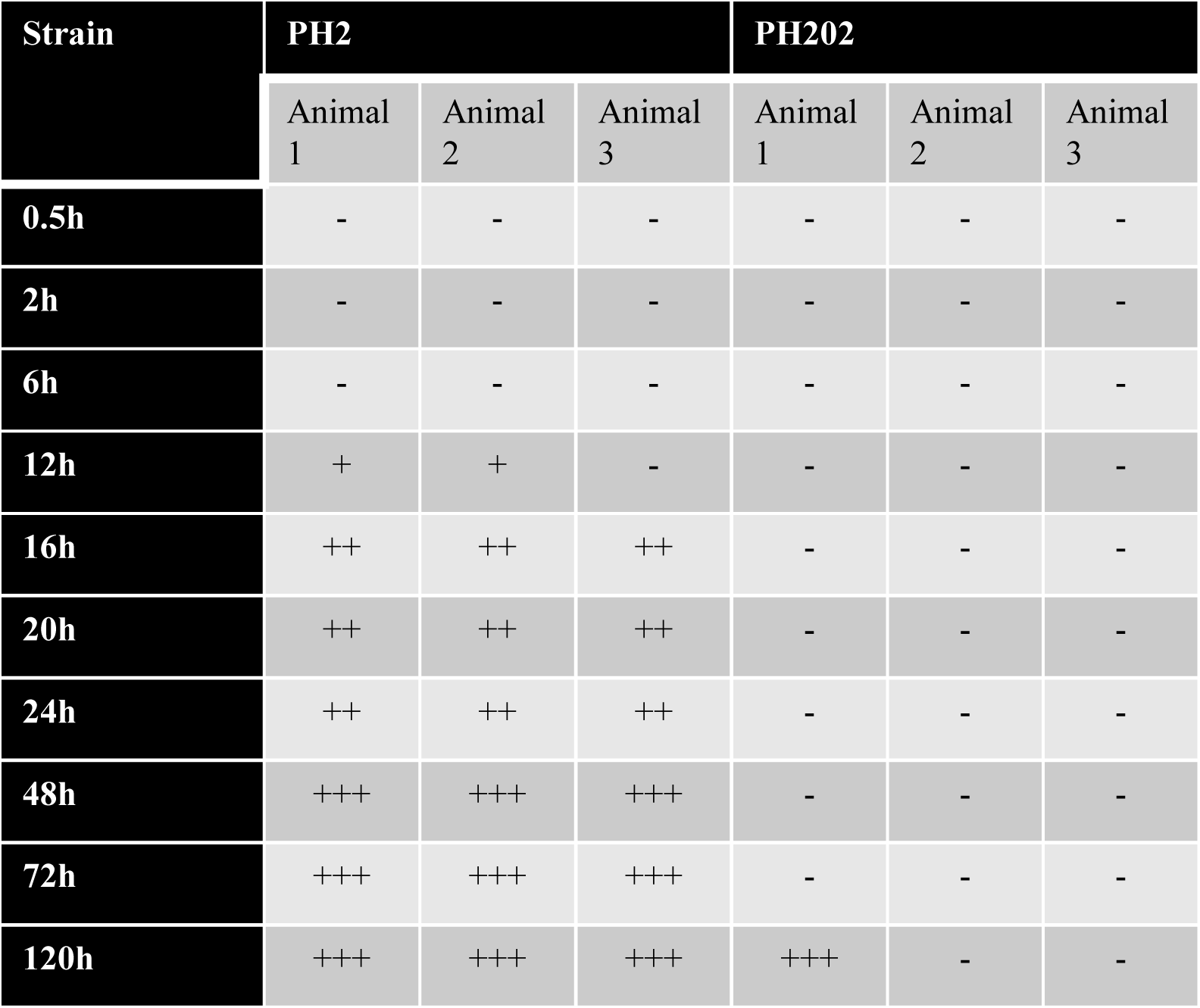
Assessment of damage to the differentiated BBEC cultures following PH2 and PH202 infection. BBEC cultures were infected apically with *M. haemolytica* strains PH2 or PH202 (2.5 × 10^7^ cfu/insert) at day 21 post-ALI. At stated time points post-infection cultures were apically washed to remove unbound bacteria, fixed and paraffin-embedded using standard histological techniques. Sections were subsequently cut, deparaffinised and H&E stained. Semi-quantitative assessment of the extent of bacterial colonisation and epithelial integrity in the histological sections was made visually. -, no sign of infection; +, low level of infection, few foci of infection present; ++, moderate level of infection, foci of infection common across the entirety of the culture; +++, high level of infection, infection present across the majority of the culture not confined to foci, cell layer showed high levels of degradation.

| Strain | PH2 |  |  | PH202 |  |  |
| --- | --- | --- | --- | --- | --- | --- |
|  | Animal 1 | Animal 2 | Animal 3 | Animal 1 | Animal 2 | Animal 3 |
| 0.5h | - | - | - | - | - | - |
| 2h | - | - | - | - | - | - |
| 6h | - | - | - | - | - | - |
| 12h | + | + | - | - | - | - |
| 16h | ++ | ++ | ++ | - | - | - |
| 20h | ++ | ++ | ++ | - | - | - |
| 24h | ++ | ++ | ++ | - | - | - |
| 48h | +++ | +++ | +++ | - | - | - |
| 72h | +++ | +++ | +++ | - | - | - |
| 120h | +++ | +++ | +++ | +++ | - | - |

### *M. haemolytica* cause intracellular infections in differentiated bovine bronchial epithelial cells

Following labelling for PH2 within infected differentiated BBEC cultures, the distribution of adherent bacteria appeared intracellular (Fig 4A [iv] & [v]). This was also observed in histological sections (Fig 4C [v]). A gentamicin protection assay was used to confirm this observation (Fig 7A). Following apical infection by PH2, a small subpopulation of gentamicin-resistant (internalised) bacteria was enumerated at 12 hpi. This population significantly increased by 24 hpi (p < 0.001, Two-way ANOVA). However, by 48 hpi this number subsequently decreased, which coincided with the reduced integrity of the epithelium. At this time point, high numbers of extracellular bacteria could be detected across the remaining tissue (Fig S4). Gentamicin-resistant (internalised) PH202 could not be detected at any time points following challenge.

**Figure 7.**
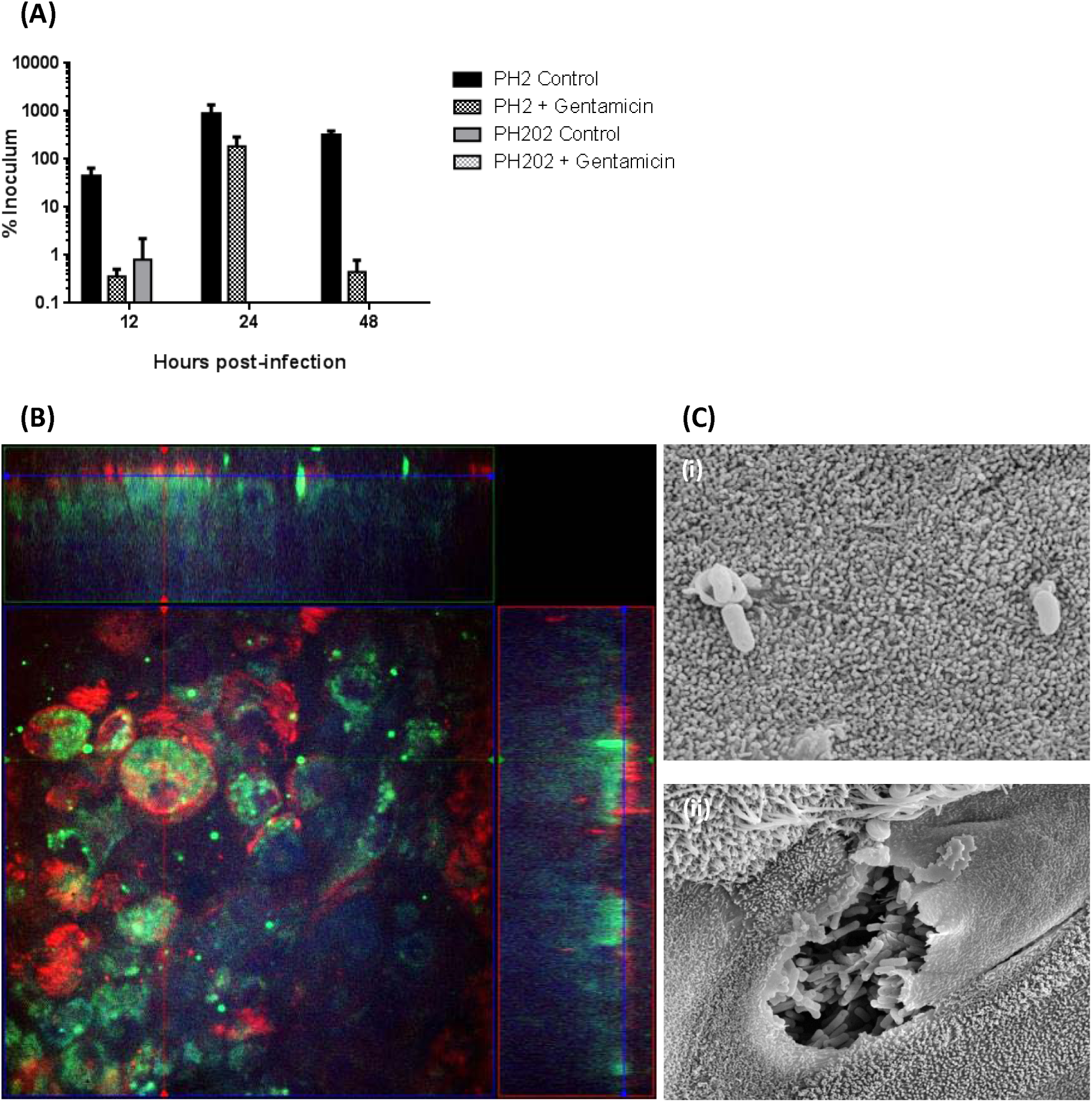
Internalisation of *M. haemolytica* in differentiated BBEC cultures. BBEC cultures were infected apically with *M. haemolytica* strains PH2 or PH202 (2.5 × 10^7^ cfu/insert) at day 21 post-ALI. At 24 hpi, cultures were apically washed to remove unbound bacteria. (A) The number of intracellular bacteria was quantified using a gentamicin protection assay, expressed as a percentage of the original inoculum. Three inserts were analysed per condition, and the data represents the mean +/- standard deviation. (B-C) BBEC cultures infected with PH2 (24 hpi) were assessed using microscopy. (B) Z-stack orthogonal representation of a BBEC culture labelled for PH2 and cilia (*M. haemolytica* - green; β-tubulin - red; nuclei – blue; x630 magnification). (C) Examination by SEM. Representative images shown of (i) invasion of a non-ciliated epithelial cell by PH2 (x10000 magnification) and (ii) an epithelial cell with apical membrane removed displaying intracellular PH2 (x5000 magnification).

Confocal microscopy confirmed the presence of intracellular *M. haemolytica* within BBECs (Fig 7B). Z-stack projections of culture 24 hpi following challenge by PH2 confirmed the presence of PH2 at near-confluent density confined within cell boundaries of both ciliated and non-ciliated epithelial cells (Fig 7B). This phenomenon was confirmed using SEM (Fig 7C). Epithelial plasma membrane projections could be observed in proximity to PH2 at the surface of a non-ciliated epithelial cell, suggesting the bacteria was being internalised via macropinocytosis (Fig 7C [i]). Internalised bacteria could also be observed at high number within epithelial cells when the apical membrane was removed (Fig 7C [ii]). This suggested *M. haemolytica* penetrated the apical surface via transcytosis, and was capable of intracellular survival within airway epithelial cells.

### *M. haemolytica* does not affect tight junction integrity in differentiated bovine bronchial epithelial cells

The integrity of tight junctions between BBECs within infected cultures was assessed following challenge by PH2 and PH202 (Fig 8). At early time points (0.5-20 hpi), tight junctions could be observed in the BBEC cultures, there was no detectable effect on the integrity of the junctional complexes due to colonisation of *M. haemolytica*. This observation was true for both the foci of infections and cells which were not colonised. Tight junctions however did appear effected at later time points following challenge by PH2 (24 hpi). Rounded epithelial cells at the foci of infection displayed reduced integrity of tight junctions at cell-to-cell borders (Fig 8A [v]). From 48-120 hpi, the epithelium was severely deteriorated and tight junctions could not be observed (Fig S7). PH202 infection however had no effect on the presence of tight junctions. These observations were confirmed by the measuring the TEER of the culture following infection (Fig 8B). Challenge by PH202 had no detectable effect on the TEER of the culture. Conversely, PH2 caused a significant reduction in TEER at 48 hpi (p < 0.001, Two-way ANOVA). This indicated that colonisation by PH2 disturbed the integrity of the tight junctions.

**Figure 8.**
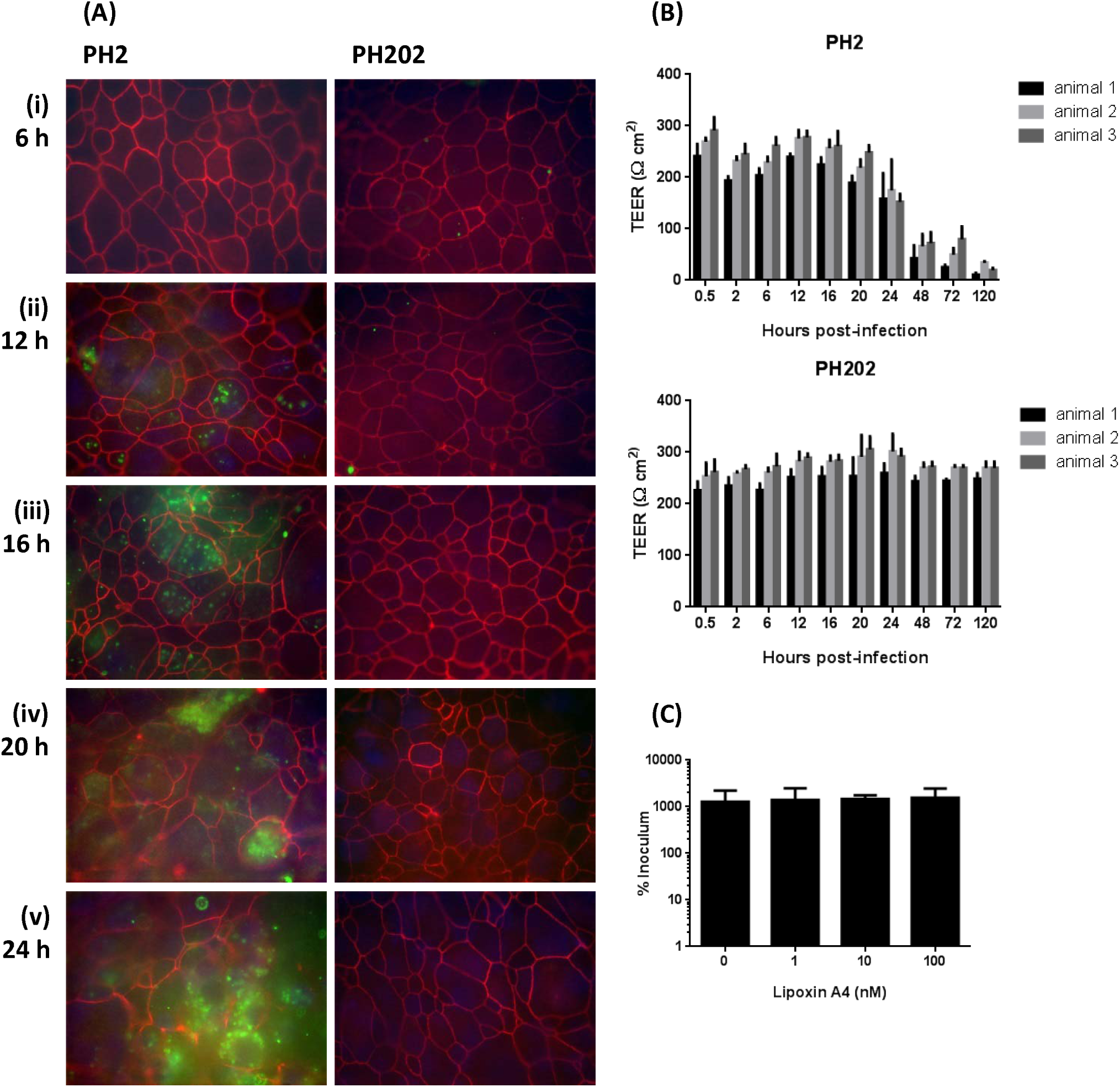
Tight junction integrity in differentiated BBEC cultures following *M. haemolytica* infection. BBEC cultures were infected apically with *M. haemolytica* strains PH2 or PH202 (2.5 × 10^7^ cfu/insert) at day 21 post-ALI. At stated time points post-infection, cultures were apically washed to remove unbound bacteria. Colonisation and tight junction integrity was subsequently assessed using (A) labelling of *M. haemolytica* and ZO-1 (*M. haemolytica* - green; ZO-1 - red; nuclei – blue; x1000 magnification). Representative images are shown of BBEC cultures at (i) 6, (ii) 12, (iii) 16, (iv) 20 and (v) 24 hpi (see Fig S7). (B) Tight-junction integrity of BBEC cultures apically infected with *M. haemolytica* was also assessed by measuring the TEER at the stated time points post-infection. (C) BBEC cultures were treated for 18 h with differing concentrations of lipoxin A4 to stabilise tight junctions prior to apical infection with *M. haemolytica* strain PH2 (2.5 × 10^7^ cfu/insert) at day 21 post-ALI. At 24 hpi, cultures were apically washed to remove unbound bacteria, and the number of adherent *M. haemolytica* quantified and expressed as a percentage of the original inoculum. For all of the above quantifications, three inserts were analysed per condition, and the data represents the mean +/- standard deviation.

This damage to junctional complexes was determined to be due to damage to the epithelium as opposed to direct targeting of tight junctions by *M. haemolytica.* Lipoxin A4 was used to stimulate tight junction formation [46]. Following treatment with lipoxin A4, TEER was increased within the BBEC culture, and labelling for ZO-1 became increasingly prominent, in a dose-dependent manner (data not shown). It was hypothesised that if PH2 was targeting tight junctions to penetrate the apical surface to colonise the culture, colonisation would be reduced following treatment. However, lipoxin A4 pre-treatment of BBECs did not affect the number of CFU adherent to the culture following challenge by PH2 24 hpi (Fig 8C).

### Serotype and host species of origin affects the capacity of *M. haemolytica* to colonise differentiated bovine bronchial epithelial cells

Differentiated BBECs were infected with eight strains of *M. haemolytica*, listed in Table 1. These strains were isolated from healthy and pneumonic cattle and sheep. Cultures were infected apically with 2.5 x 10^7^ cfu/insert, and colonisation was quantified at 2, 24 and 72 hpi (Fig 9A [i] & S8). The number of CFU present in the apical wash was also enumerated (Fig 9 [ii] 7 S8). At 2 hpi, there was no significant difference in the adherence efficiency between all eight strains (Two-way ANOVA). There was little evidence of bacteria present on the apical surface as observed using SEM at 2 hpi (Fig S10 & S11). By 24 hpi however, both the virulent A1 and A6 strains isolated from pneumonic cattle (PH2 and PH376 respectively) and virulent A2 strains isolated from pneumonic sheep (PH278 and PH372) were capable of successfully colonising the BBEC cultures. This colonisation was observed as foci of infection below the apical surface, as described for PH2 (Fig 10A). The apical surfaces surrounding the foci were largely devoid of adherent bacteria (Fig 10B). Adherence of virulent bovine strains to the BBEC was approximately 10-fold higher in comparison to virulent ovine strains. This was reflected in a higher number of foci of infection present after challenge with PH2 and PH376. At 72 hpi, significant deterioration was observed in epithelia infected by all virulent strains, as displayed within histological sections and SEM (Fig 10). This was reflected in a reduction in the TEER (Fig 9B). A semi-quantitative scoring of this damage using histological sections was performed (Table 3). Deterioration following infection by virulent ovine strains, particularly PH372 was less prominent in comparison to the bovine strains, which appeared more invasive to BBEC cultures.

**Figure 9.**
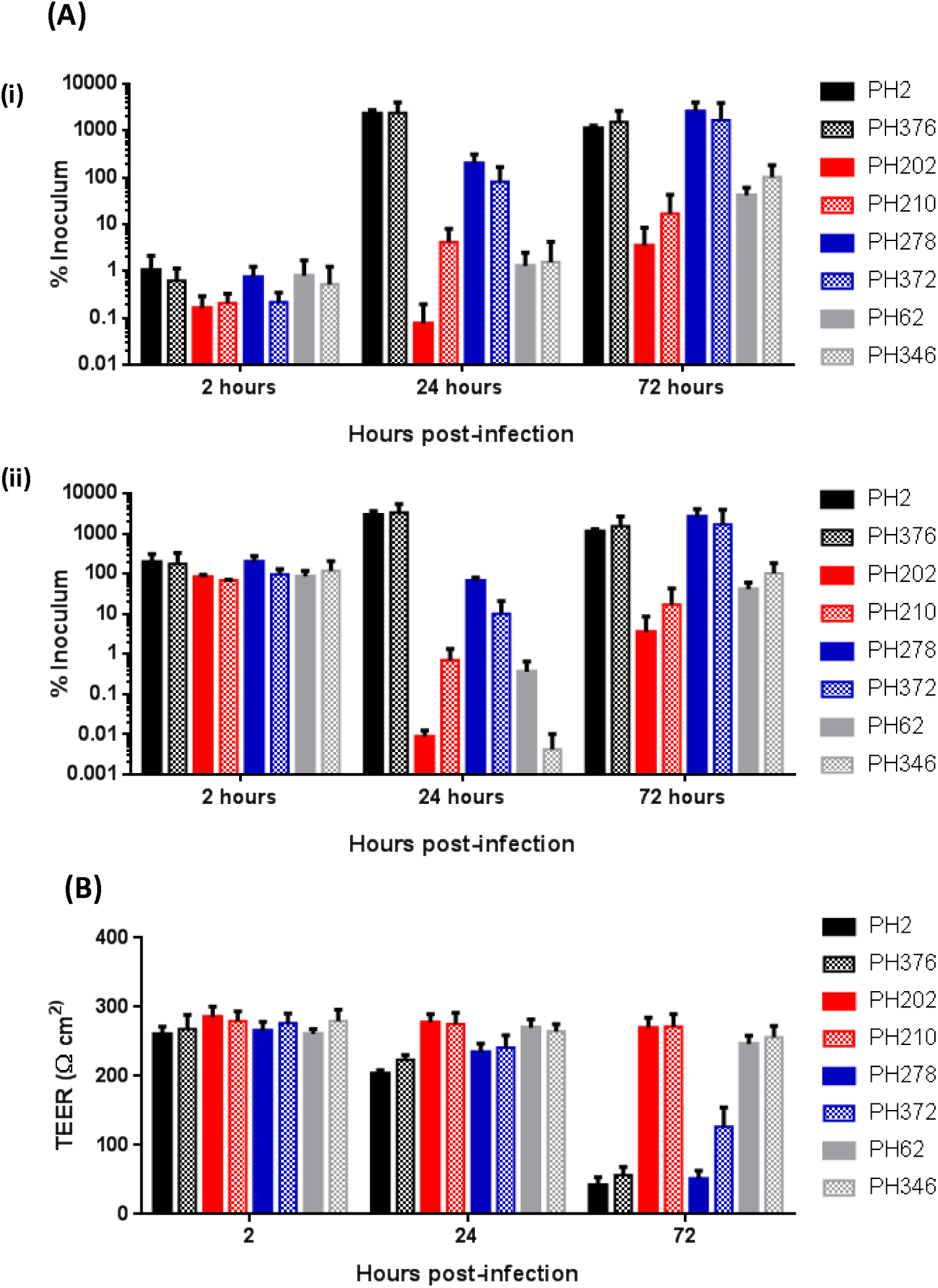
Quantification of adhesion of *M. haemolytica* strains to differentiated BBEC cultures. BBEC cultures were infected apically with eight strains of *M. haemolytica* (2.5 × 10^7^ cfu/insert) at day 21 post-ALI. (A) At stated time points post-infection, cultures were apically washed to remove unbound bacteria, and colonisation assessed. Quantification of (i) the number of adherent *M. haemolytica* and (ii) *M. haemolytica* present in the apical wash, as expressed as a percentage of the original inoculum. (B) Tight-junction integrity of BBEC cultures apically infected with eight strains *M. haemolytica* was also assessed by measuring the TEER at the stated time points post-infection. For all of the above quantifications, three inserts were analysed per condition, and the data represents the mean +/- standard deviation from cultures derived from three different animals (see Fig S8).

Neither the commensal bovine (PH202 and PH210, A2 serotype) or ovine strains (PH62 and PH346, A12 serotype) displayed a significant increases in adherence efficiency between 2 to 24 hpi (Two-way ANOVA; Fig 9). There was a slight increase in CFU present in the culture for all commensal strains by 72 hpi; however the number of bacteria within the culture was significantly lower in comparison to all strains isolated from pneumonic animals. Visually, BBEC cultures infected by commensal strains did not present overt evidence of colonisation or damage to the epithelium (Fig 10), and the TEER was not affected (Fig 9B). Foci of infection could be observed in individual cultures for PH210, PH62 and PH346 (Table 3). However these were in isolated incidence and were present at a much lower number in comparison to virulent strains. Such foci were located towards the border regions of the foci, at which the epithelium can present evidence of damage. This data suggests that virulent strains were capable of successfully colonising differentiated BBEC following apical infection. Conversely, commensal strains isolated from the nasopharynx of healthy animals were not capable of successfully colonising the model to a high degree.

**Figure 10.**
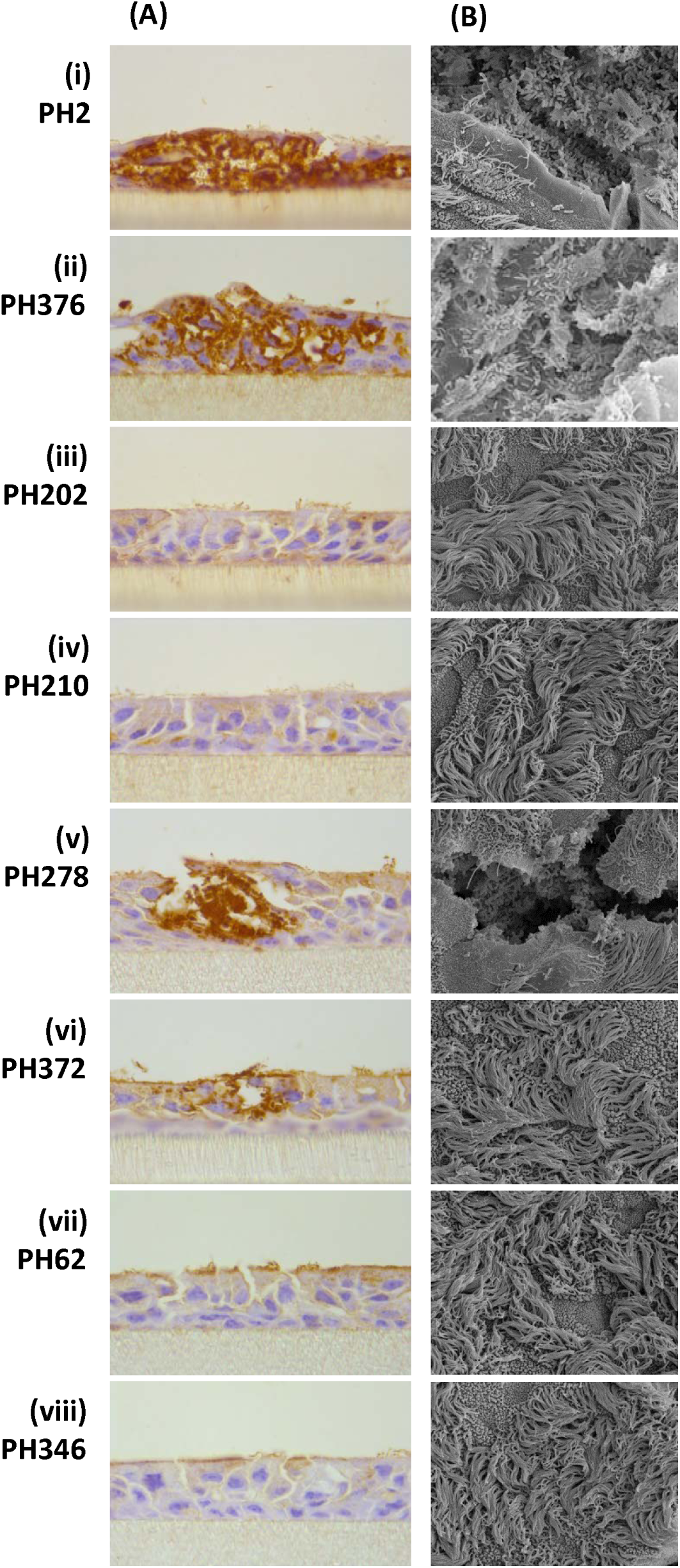
Colonisation of *M. haemolytica* strains to differentiated BBEC cultures. BBEC cultures were infected apically with eight strains of *M. haemolytica* (2.5 × 10^7^ cfu/insert) at day 21 post-ALI. At 24 hpi, cultures were apically washed to remove unbound bacteria, and fixed. Colonisation of *M. haemolytica* was subsequently assessed using (A) immunohistochemical-labelling of *M. haemolytica* in histological sections (OmpA-labelled *M. haemolytica* stained brown; x1000 magnification) and (B) examination by SEM (x2500 magnification). Representative images are shown of BBEC cultures infected with (i) PH2, (ii) PH376, (iii) PH202, (iv) PH210, (v) PH278, (vi) PH376, (vii) PH62 and (viii) PH346 (see Fig S9, S10 and S11).

**Table 3.** Assessment of damage to the differentiated BBEC cultures following *M. haemolytica* infection. BBEC cultures were infected apically with eight strains of *M. haemolytica* (2.5 × 10^7^ cfu/insert) at day 21 post-ALI. At stated time points post-infection cultures were apically washed to remove unbound bacteria, fixed and paraffin-embedded using standard histological techniques. Sections were subsequently cut, deparaffinised and H&E stained (see Fig S12). Semi-quantitative assessment of the extent of bacterial colonisation and epithelial integrity in the histological sections was made visually. -, no sign of infection; +, low level of infection, few foci of infection present; ++, moderate level of infection, foci of infection common across the entirety of the culture; +++, high level of infection, infection present across the majority of the culture not confined to foci, cell layer showed high levels of degradation.

| Strain | 2 hours |  |  | 24 hours |  |  | 72 hours |  |  |
| --- | --- | --- | --- | --- | --- | --- | --- | --- | --- |
|  | Animal 1 | Animal 2 | Animal 3 | Animal 1 | Animal 2 | Animal 3 | Animal 1 | Animal 2 | Animal 3 |
| PH2 | - | - | - | ++ | ++ | ++ | +++ | +++ | +++ |
| PH376 | - | - | - | ++ | ++ | ++ | +++ | +++ | +++ |
| PH202 | - | - | - | - | - | - | - | - | - |
| PH210 | - | - | - | + | - | - | - | - | - |
| PH278 | - | - | - | ++ | ++ | + | +++ | +++ | ++ |
| PH372 | - | - | - | + | + | + | ++ | + | ++ |
| PH62 | - | - | - | - | + | - | + | - | - |
| PH346 | - | - | - | + | + | - | - | - | - |

## Discussion

The aim of the study was to investigate the interaction between pathogenic and virulent strains of *M. haemolytica* with the bovine airway epithelium using a differentiated cell model, in order to have a better understanding of the events of BRD. Differentiated airway epithelial models have been utilised previously to study a number of bacterial pathogens, including *Pseudomonas aeruginosa* [47-49], *Haemophilus influenzae* [50-52], *Neisseria meningitidis* [53] and *Mycoplasma pneumoniae* [38, 41]. To our knowledge, this investigation is the first to utilise a differentiated cell model to study the interaction of *M. haemolytica* with the bovine airway epithelium. This has allowed the adherence, colonisation and traversal of the epithelium by *M. haemolytica* to be studied in a model which displayed the characteristic defence mechanisms associated with the respiratory tract. The model provides an alternative to animal models, which are costly and time-intensive, and are contrary to the three R’s principles.

Submerged BBECs have routinely been utilised to study the adherence of *M. haemolytica* [30, 54-56]. However, submerged epithelial cultures are undifferentiated [57], and as such do not replicate the complexity of the airway epithelium [58]. Bovine bronchial epithelial cells can be stimulated to differentiate into a more representative model of the *in vivo* microenvironment through exposure to the atmosphere [36, 37, 59]. The methodology for differentiation of bovine airway epithelial cells have been fully optimised [44] and the model well-characterised [45]. The differentiated BBEC model has been shown to replicate the hallmark defences of the respiratory tract, including active mucocilary clearance. These mechanisms actively prevent colonisation of invading pathogens, and are important considerations when modelling bacterial interactions within the airway. Tight junctions present in the differentiated BBEC also allowed for the identification of invasion of the pathogen through the apical barrier (Fig 4C) into the sub-apical epithelium. Our model therefore allows for these defence mechanisms to be considered when assessing colonisation of *M. haemolytica*, and as such provides an excellent model to characterise *M. haemolytica-*host interaction in the bovine airway epithelium.

A direct comparison of the ability of *M. haemolytica* to colonise differentiated and undifferentiated BBEC cultures was made in this investigation (Fig 1 & Fig 3). Initial adherence of *M. haemolytica* to BBEC was comparable between the two models. As undifferentiated BBEC cultures do not possess cilia, it was inferred that adhesion is not specific to either ciliated or unciliated cells. This was confirmed using immunofluorescence and SEM (Fig 4). In both models PH2, isolated from lung of pneumonic cattle, was capable of heavily colonising cells by 24 hpi. Adherence at 24 hpi was 10-fold higher in the differentiated model in comparison to the undifferentiated model. This may be due to the increased thickness of the epithelium in the differentiated model, where a columnar, pseudostratified morphology was formed, stereotypical for the airway epithelium (Fig 2), as opposed to a two-dimensional monolayer. This provided a larger 3-D architecture for the bacteria to adhere, highlighting the importance of cell differentiation when investigating epithelial colonisation.

The differentiated BBEC was initially infected with two strains of *M. haemolytica*, PH2, a bovine isolate from the lung of a pneumonic animal, and PH202, a bovine isolate from the nasopharynx of a healthy animal (Fig 3). PH2 is an A1 serotype strain, which is the major cause of pneumonic pasteurellosis in cattle [60, 61]. PH202 is an A2 serotype, a predominately non-pathogenic serotype present as a commensal species in cattle [29, 61]. A1 and A2 are the most prevalent serotypes worldwide [29, 60]. Both strains have previously been shown to adhere to undifferentiated ovine bronchial epithelial cells [62]. Initial adherence of both PH2 and PH202 to differentiated BBEC cultures was low (Fig 3). At early time points (0.5-6 hpi), bacteria associated with the culture were distributed at a low density across the apical surface (Figs 4A & 5A). However, following further incubation PH2 became increasingly abundant. Penetration of PH2 below the apical surface of the culture was identified by 12 hpi; suggesting *M. haemolytica* A1 is capable of traversing epithelial tight junctions (Fig 4C). This coincided with a significant increase in the number of adherent bacteria (Fig 3). At the foci of infection, PH2 replicated to a high density below the apical surface (Fig 4). Such foci have previously been described for *H. influenza* [51] and *M. pneumoniae* [41]. The number of foci increased with exposure time, as did the number of PH2 present in the systems which were not closely associated with tissue (Fig 3). This provided evidence that the bacteria disseminated from the foci to re-infecting other regions of the culture. From 48 hpi onwards, the majority of the epithelium was heavily infected, and distinct foci no longer observed.

Conversely, A2 commensal strain PH202 was not capable of colonising the differentiated BBEC model. Although initial adherence of the bacterium could be detected at early time points post-infection (Fig 3), there was no evidence of PH202 penetrating the apical surface (Fig 5C). In tissue derived from two of the three animals, viable PH202 could not be recovered from the culture at the end point of the infection. This finding indicated that the differentiated model actively prevented colonisation by the bacteria. This may be due to mucociliary clearance which actively removes invading pathogens [63, 64]. Alternatively, BBECs are known to produce anti-bacterial peptides, such as tracheal antimicrobial peptide (TAP), in response to bacterial products including LPS [65-67]. This peptide has shown to be bactericidal against *M. haemolytica* [68]. The variation observed in the ability of the A1 and A2 bacteria to adhere to the culture may provide insight into the selective explosion of the A1 population over A2 strains, resulting in infection in the lower respiratory tract during pneumonic pasteurellosis [27].

Following extended co-culture of differentiated BBECs and PH2 (48-120 hpi), there was significant evidence of damage present in airway epithelial cells. By 24 hpi, there was cell rounding and detachment in BBECs heavily colonised by *M. haemolytica* (Fig 6). This phenomenon was more pronounced at 48 hpi, where a large number of cells were readily detached from culture following apical washing (Fig S6). This response was only observed following PH202 infection in tissue derived from one of three animals at 120 hpi (Table 2). Similar cytopathic effects in epithelial cells have been observed in response to other bacterial pathogens, including *P. aeruginosa* [69] and *Klebsiella pneumoniae* [70]. This may mimic events *in vivo*. Clinical signs of *M. haemolytica* include pulmonary lesions, and necrosis and desquamation can also be observed at the bronchial epithelium [29]. The cause of the induced cell death in the model is unknown. Major *Mannheimia* virulence factors lipopolysaccharide (LPS) and leukotoxin have been shown to not cause necrosis or apoptosis in bovine pulmonary epithelial cells [71]. Epithelial cell death observed in the system may however be due to the innate immune response following bacterial infection, resulting in the sloughing off of infected cells [72].

In this study we presented evidence that *M. haemolytica* A1 strain PH2 may be internalised by airway epithelial cells following infection (Fig 7). Internalisation of *M. haemolytica* A1 by BBECs has not been previously reported. A small subpopulation of PH2 was identified to be resistant following treatment with gentamicin, indicative of the presence of intracellular bacteria (Fig 7A). This was confirmed using a number of microscopy techniques (Figs 7B and 7C). Within BBECs, intracellular PH2 could be detected from 12 hpi. This number appeared to peak at 24 hpi (Fig 7A). Electron microscopy suggests that internalisation of PH2 in epithelial cells may be occurring through macropinocytosis (Fig 7C). This has previously been observed for *H. influenzae* [51]*. M. haemolytica* was present at high density within cell boundaries (Fig 7B), suggesting that once internalised PH2 is capable of survival and replication within host cells. *M. haemolytica* A1 may invade BBECs in order to gain access to sub-epithelial spaces thorough transcytosis of the epithelium. An intracellular phase during infection may also allow for persistence or evasion of aspects of host immunity, such as humoral and complement-attack or mucociliary clearance [73].

Tight junctions create a physiochemical barrier to prevent the invasion of pathogens from the lumen of the airway to the interstitial compartment [74, 75]. However, several bacteria are capable of disrupting the junctional complexes during paracytosis [76, 77]. This was not observed following colonisation of either PH2 or PH202 (Fig 8). Tight junctions were present following challenge by *M. haemolytica*, with no detectable reduction in integrity by 24 hpi (Fig 8A). This was confirmed by measuring the TEER of the culture (Fig 8B). There was however a detectable decrease in the number of tight junctions present by 48 hpi with PH2. However this was due to significant damage to the epithelium following infection, particularly at the apical surface (Fig S6). The addition of lipoxin A4, which stimulates expression of ZO-1, has previously been shown to prevent the invasion and transmigration of *P. aeruginosa* [78]. However stimulation of BBEC cultures did not prevent colonisation by PH2, despite enhancing the integrity of tight junctions (Fig 8C), further suggesting that tight junctions are not targeted by *M. haemolytica*. It is hypothesised therefore that transmigration of the bacterium through the apical surface of the epithelium occurred via transcytosis, but not through paracellular transport.

The pattern of infection of PH2 was replicated using a second bovine A6 isolate PH376. Both strains colonised the BBEC cultures to a comparable degree (Fig 9A), forming morphologically similar infection foci (Fig 10) beneath the apical surface at 24 hpi. Significant damage to the tissue was detected at 72 hpi with both strains (Fig S12). A2 bovine isolate PH210, in agreement with PH202, was not capable of forming foci of infection stereotypical of A1 strains (Fig 10). The model was further challenged with four ovine isolates (Fig 9 & 10). Two strains (PH278 and PH372) were representative of A2 ovine strains, the major causative agent of pneumonia in sheep [29, 60]. As with the bovine isolates, the virulent strains were capable of colonising the model to form foci of infection. The A2 bovine isolates behaved differently from ovine A2 isolates when co-cultured with the model (Fig 9). This is to be expected as the outer-membrane protein profiles differ between the two groups [21]. Cultures were also infected with two A12 strains (PH62 and PH346), which are traditionally not associated with disease. The commensal A12 strains failed to colonise the cultures to a similar degree (Fig 9). Adherence to bovine airway epithelial cells by virulent ovine isolates was approximately 10-fold lower in comparison to virulent bovine isolates (Fig 9). This suggests that host specificity in *M. haemolytica* strains was partly dependent on specific cell-surface structures present on differentiated BBECs. This difference in pathogenesis between serotypes is likely due to variation in the LPS profile [21, 22], outer-membrane proteins [21] and allelic variation in a number of virulence genes such as *lktA* [23], *ompA* [24]. In conclusion, variation in disease pathogenesis *in vivo* due to serotype and host specificity is reflected in the degree of colonisation within the differentiated BBEC culture.

In this study we have characterised the host-pathogen interactions between BBECs grown at an ALI with various serotypes of *M. haemolytica*. The model used to investigate the infection *in vitro* has been shown to be highly representative of the *in vivo* epithelium, providing insight into the pathogenesis of *M. haemolytica* during pneumonic pasteurellosis in the context of host defence mechanisms, such as tight junctions and mucociliary clearance. *M. haemolytica* A1 was capable of highly colonising the model, causing extensive damage to the host epithelium. This occurred at foci of infection, below the apical surface of the tissue. Tight junctions in the epithelium were bypassed using transcytosis, but not paracytosis. *M. haemolytica* A2 was not capable of replicating this colonisation. This may account for the occurrence of lower respiratory tract infection following the shift from A2 to A1 population in cattle prior to onset of pneumonic pasteurellosis. The BBEC model was further challenged using a panel of isolates, and the degree of pathogenesis was dependent on both serotype and host species. This investigation provides the first insight into the route of infection of *M. haemolytica* in a differentiated model of the bovine airway epithelium.

## Acknowledgments

We thank Ms Margaret Mullin and Ms Lynne Stevenson (both University of Glasgow) for assistance with electron microscopy and histology, respectively.

---

**Supplementary Figure 1.**
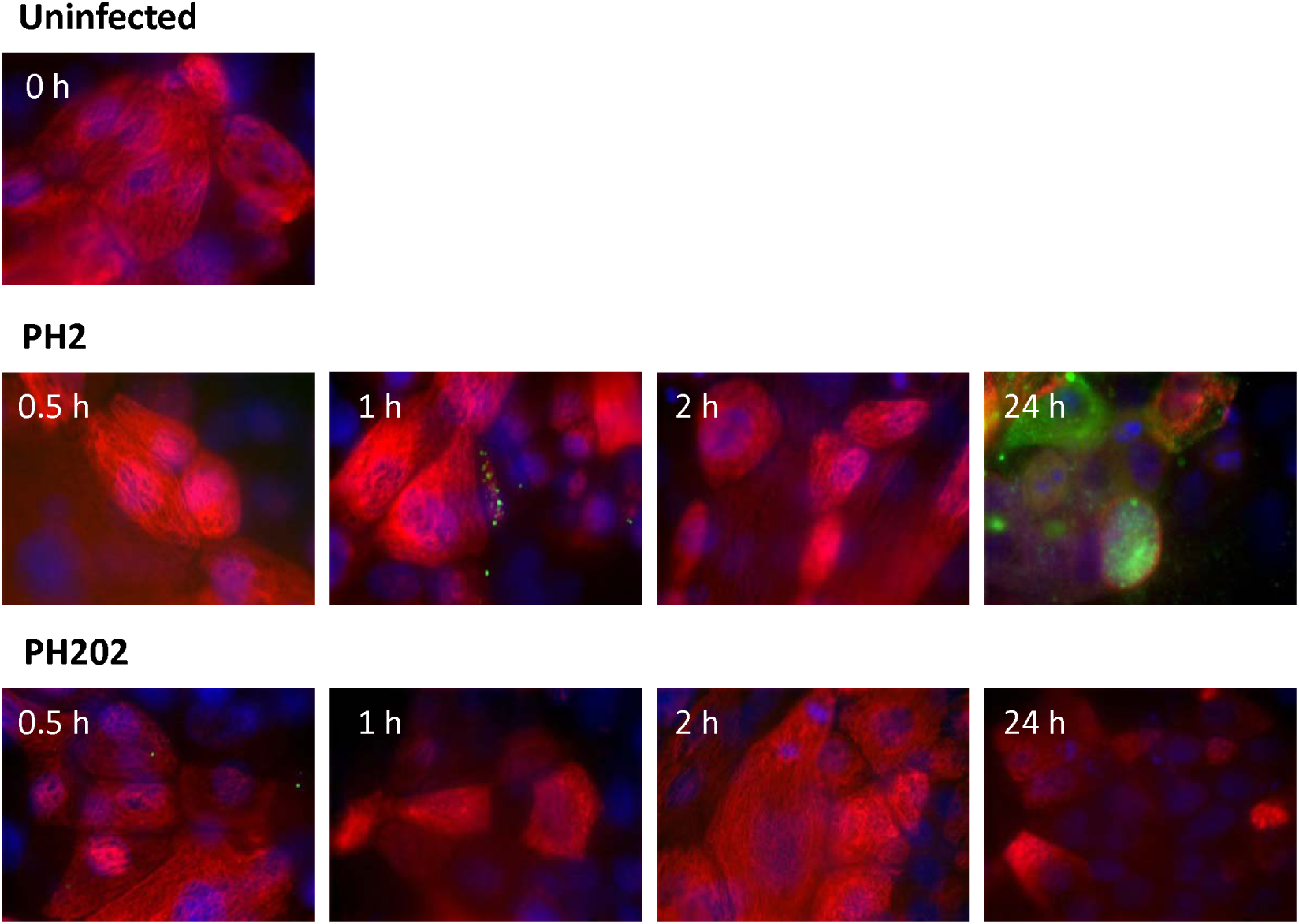
Immunofluorescent-labelling of *M. haemolytica* and β-tubulin following infection of undifferentiated BBEC cultures. BBEC cultures were infected apically with *M. haemolytica* strains PH2 or PH202 (2.5 × 10^7^ cfu/insert) at day 0 post-ALI. At 0.5, 1, 2 and 24 hpi, cultures were apically washed to remove unbound bacteria, and fixed. Colonisation of PH2 and PH202 was subsequently assessed using immunofluorescence labelling of *M. haemolytica* and β-tubulin (*M. haemolytica* - green; β-tubulin - red; nuclei – blue; x1000 magnification).

**Supplementary Figure 2.**
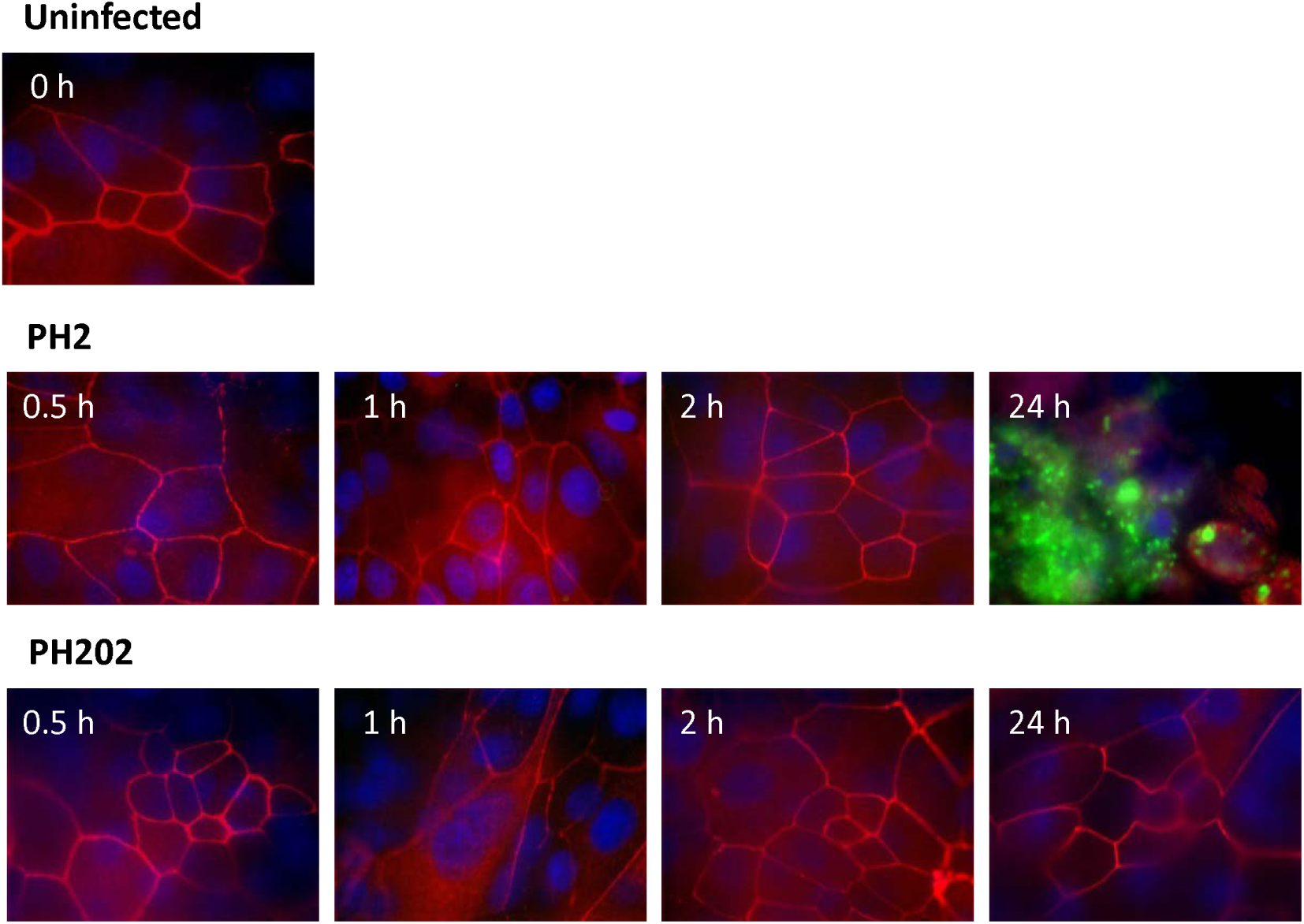
Immunofluorescent-labelling of *M. haemolytica* and ZO-1 following infection of undifferentiated BBEC cultures. BBEC cultures were infected apically with *M. haemolytica* strains PH2 or PH202 (2.5 × 10^7^ cfu/insert) at day 0 post-ALI. At 0.5, 1, 2 and 24 hpi, cultures were apically washed to remove unbound bacteria, and fixed. Colonisation of PH2 and PH202 was subsequently assessed using immunofluorescence labelling of *M. haemolytica* and tight junctions (*M. haemolytica* - green; ZO-1 - red; nuclei – blue; x1000 magnification).

**Supplementary Figure 3.**
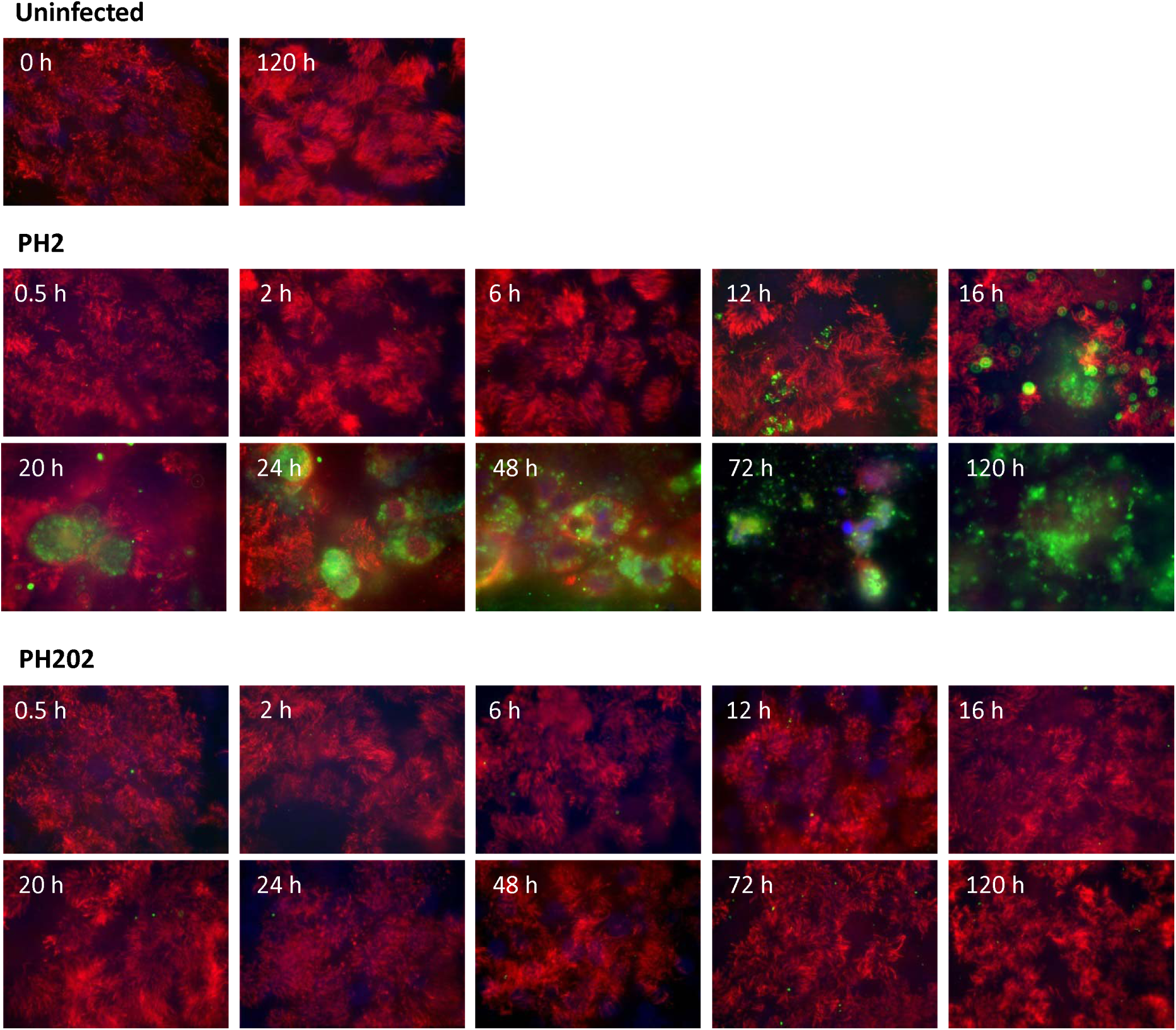
Immunofluorescent-labelling of PH2 or PH202 and β-tubulin following infection of differentiated BBEC cultures. BBEC cultures were infected apically with *M. haemolytica* strains PH2 or PH202 (2.5 × 10^7^ cfu/insert) at day 21 post-ALI. At stated time points post-infection, cultures were apically washed to remove unbound bacteria, and fixed. Colonisation of PH2 and PH202 was subsequently assessed using immunofluorescence labelling of *M. haemolytica* and cilia (*M. haemolytica* - green; β-tubulin - red; nuclei – blue; x1000 magnification).

**Supplementary Figure 4.**
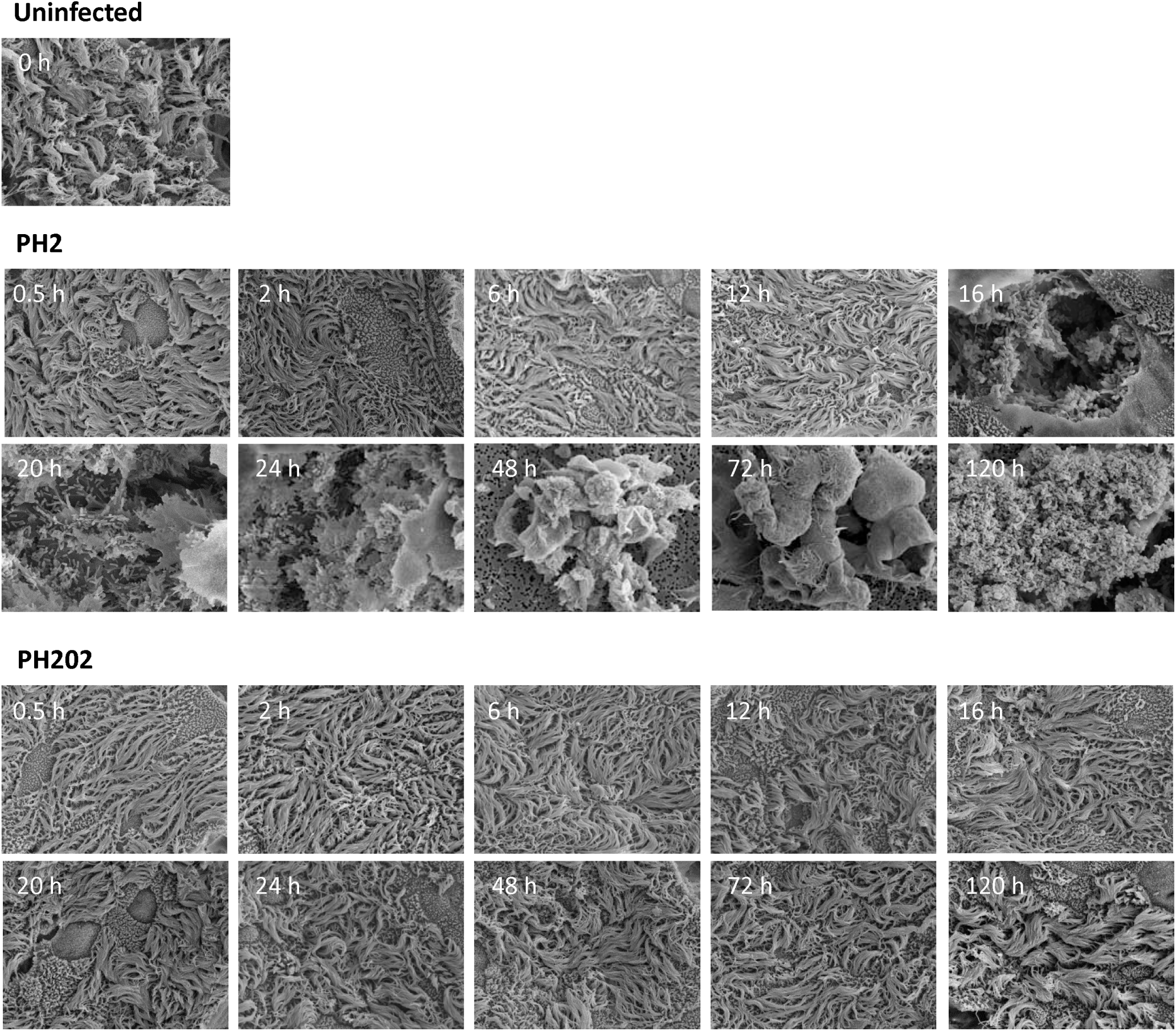
SEM examination of PH2 or PH202 infection of differentiated BBEC cultures. BBEC cultures were infected apically with *M. haemolytica* strains PH2 or PH202 (2.5 × 10^7^ cfu/insert) at day 21 post-ALI. At stated time points post-infection, cultures were apically washed to remove unbound bacteria, fixed and examination by SEM (x2500 magnification).

**Supplementary Figure 5.**
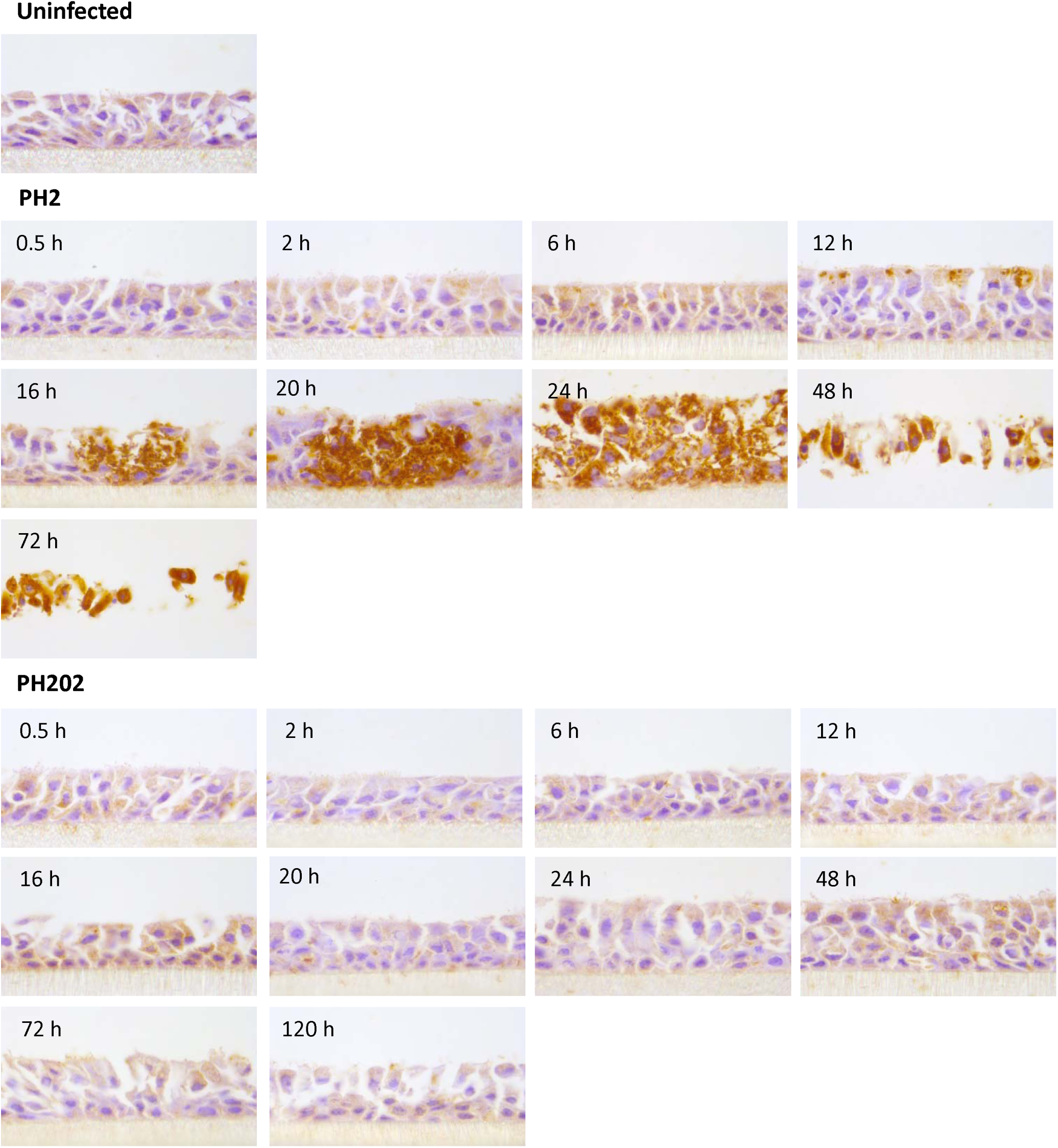
Immunohistochemical-labelling of PH2 or PH202 in histological sections following infection of differentiated BBEC cultures. BBEC cultures were infected apically with *M. haemolytica* strains PH2 or PH202 (2.5 × 10^7^ cfu/insert) at day 21 post-ALI. At stated time points post-infection, cultures were apically washed to remove unbound bacteria, fixed and paraffin-embedded using standard histological techniques. Sections were subsequently cut, deparaffinised and immunohistochemistry labelling of *M. haemolytica* was performed using an anti-OmpA antibody (OmpA-labelled *M. haemolytica* stained brown; x1000 magnification). For PH2 120 hpi, the tissue layer was too damaged to recover following antigen retrieval.

**Supplementary Figure 6.**
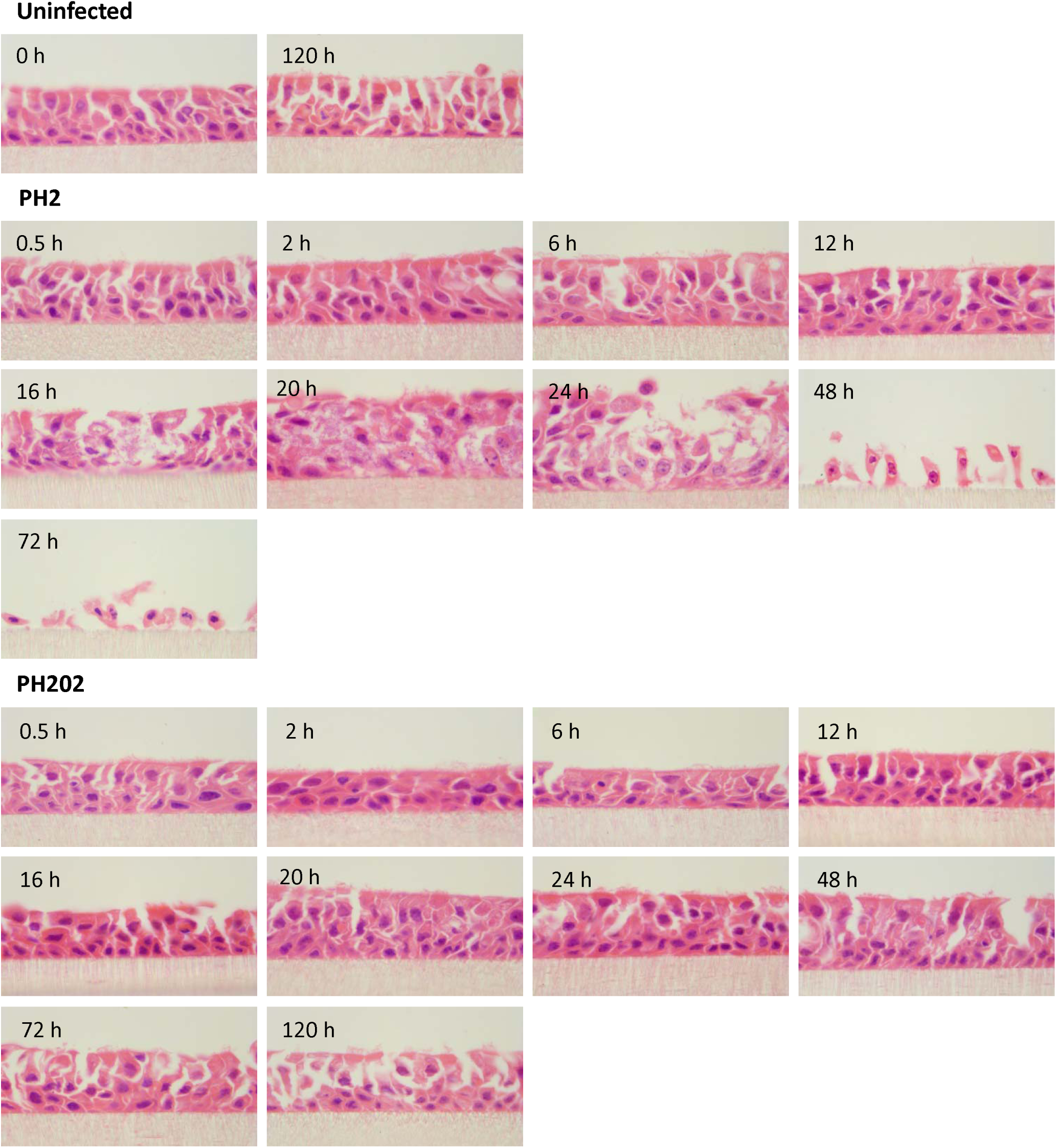
H&E staining of histological sections following PH2 or PH202 infection of differentiated BBEC cultures. BBEC cultures were infected apically with *M. haemolytica* strains PH2 or PH202 (2.5 × 10^7^ cfu/insert) at day 21 post-ALI. At stated time points post-infection, cultures were apically washed to remove unbound bacteria, fixed and paraffin-embedded using standard histological techniques. Sections were subsequently cut, deparaffinised and H&E staining was performed (x1000 magnification).

**Supplementary Figure 7.**
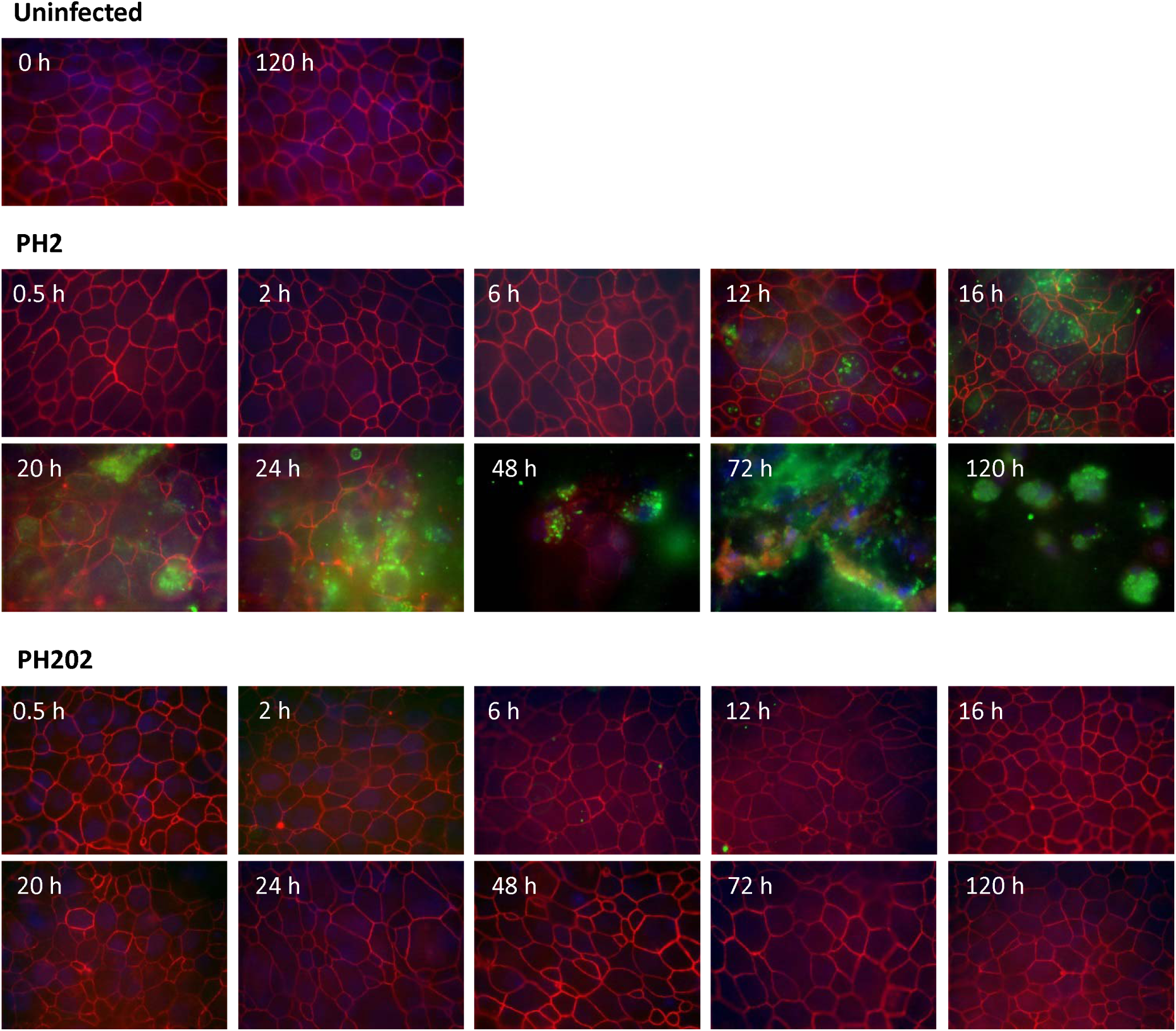
Immunofluorescent-labelling of PH2 or PH202 and ZO-1 following infection of differentiated BBEC cultures. BBEC cultures were infected apically with *M. haemolytica* strains PH2 or PH202 (2.5 × 10^7^ cfu/insert) at day 21 post-ALI. At stated time points post-infection, cultures were apically washed to remove unbound bacteria, and fixed. Colonisation of PH2 and PH202 and tight junction integrity was subsequently assessed using immunofluorescence labelling of *M. haemolytica* and tight junctions (*M. haemolytica* - green; ZO-1 - red; nuclei – blue; x1000 magnification).

**Supplementary Figure 8.**
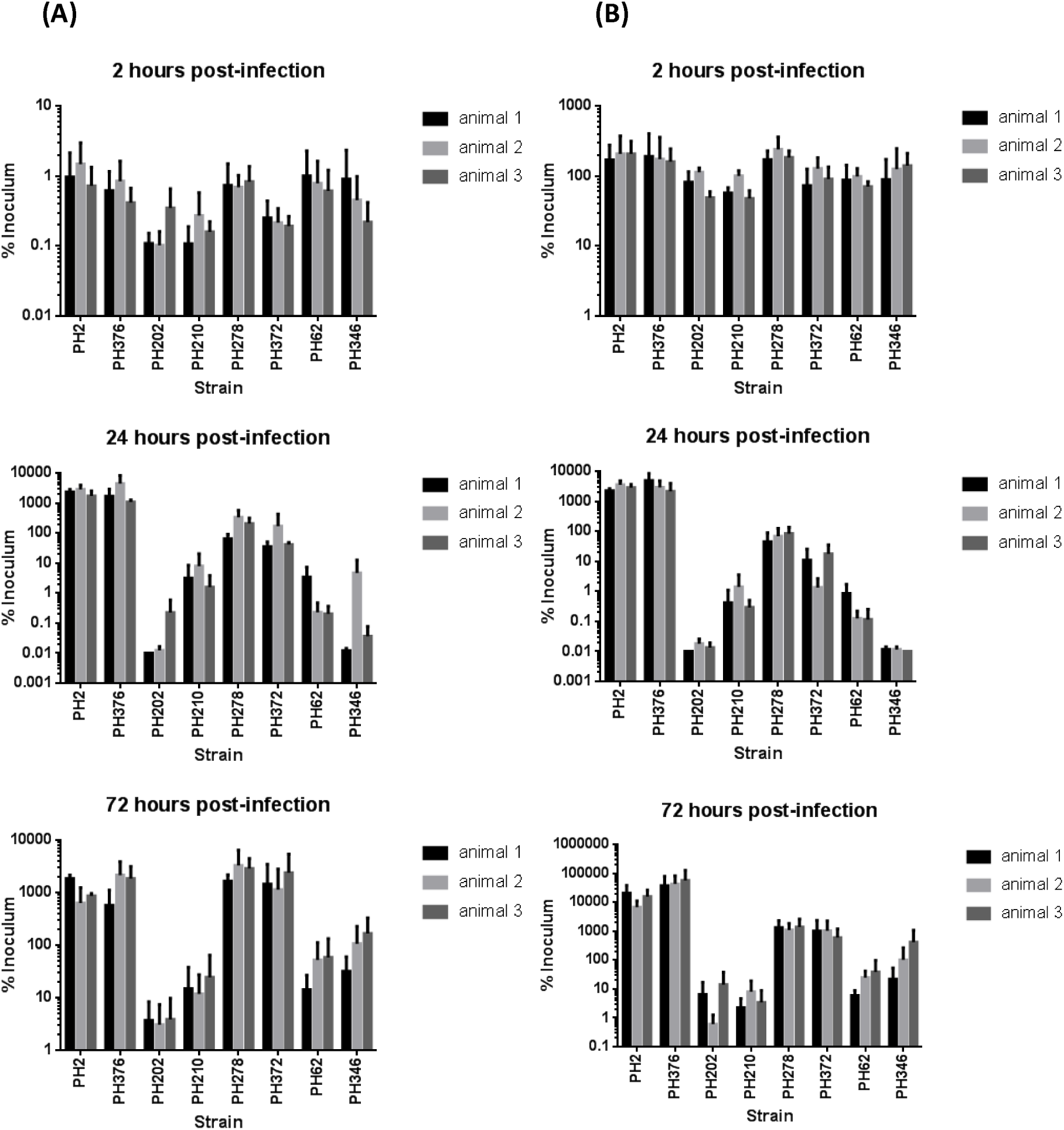
Quantification of adhesion of *M. haemolytica* strains to differentiated BBEC cultures. BBEC cultures were infected apically with eight strains of *M. haemolytica* (2.5 × 10^7^ cfu/insert) at day 21 post-ALI. At 2, 24 or 72 hpi, cultures were apically washed to remove unbound bacteria, and colonisation assessed. Quantification of (A) the number of adherent *M. haemolytica* and (B) *M. haemolytica* present in the apical wash, as expressed as a percentage of the original inoculum. Three inserts were analysed per time point, and the data represents the mean +/− standard deviation.

**Supplementary Figure 9.**
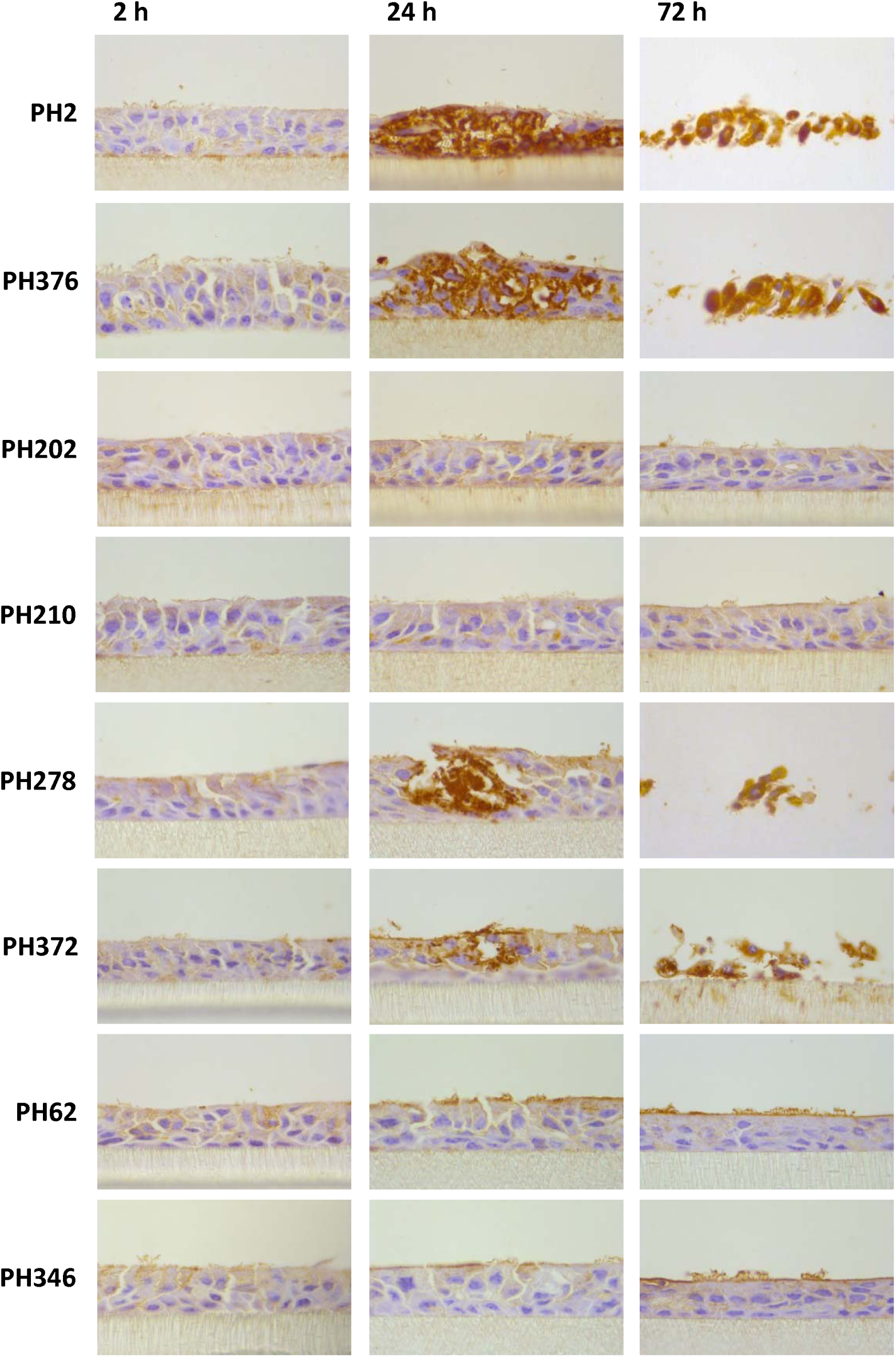
Immunohistochemical-labelling of *M. haemolytica* strains in histological sections following infection of differentiated BBEC cultures. BBEC cultures were infected apically with eight strains of *M. haemolytica* (2.5 × 10^7^ cfu/insert) at day 21 post-ALI. At 2, 24 or 72 hpi, cultures were apically washed to remove unbound bacteria, fixed and paraffin-embedded using standard histological techniques. Sections were subsequently cut, deparaffinised and immunohistochemistry labelling of *M. haemolytica* was performed using an anti-OmpA antibody (OmpA-labelled *M. haemolytica* stained brown; x1000 magnification).

**Supplementary Figure 10.**
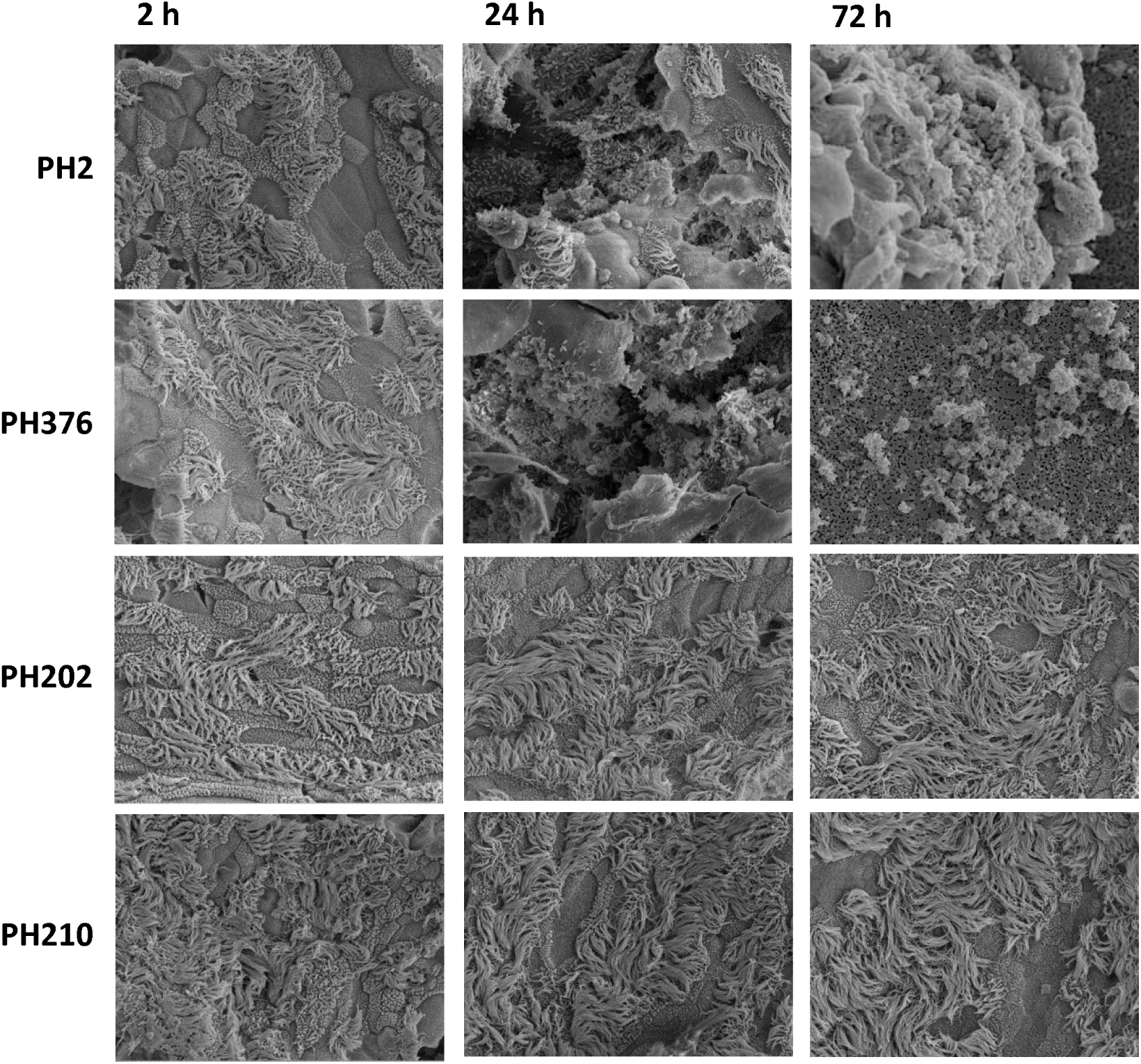
SEM examination of differentiated BBEC culture infected with bovine *M. haemolytica* isolates. BBEC cultures were infected apically with *M. haemolytica* strains PH2, PH376, PH202 or PH210 (2.5 × 10^7^ cfu/insert) at day 21 post-ALI. At 2, 24 or 72 hpi, cultures were apically washed to remove unbound bacteria, fixed and examination by SEM (x2500 magnification).

**Supplementary Figure 11.**
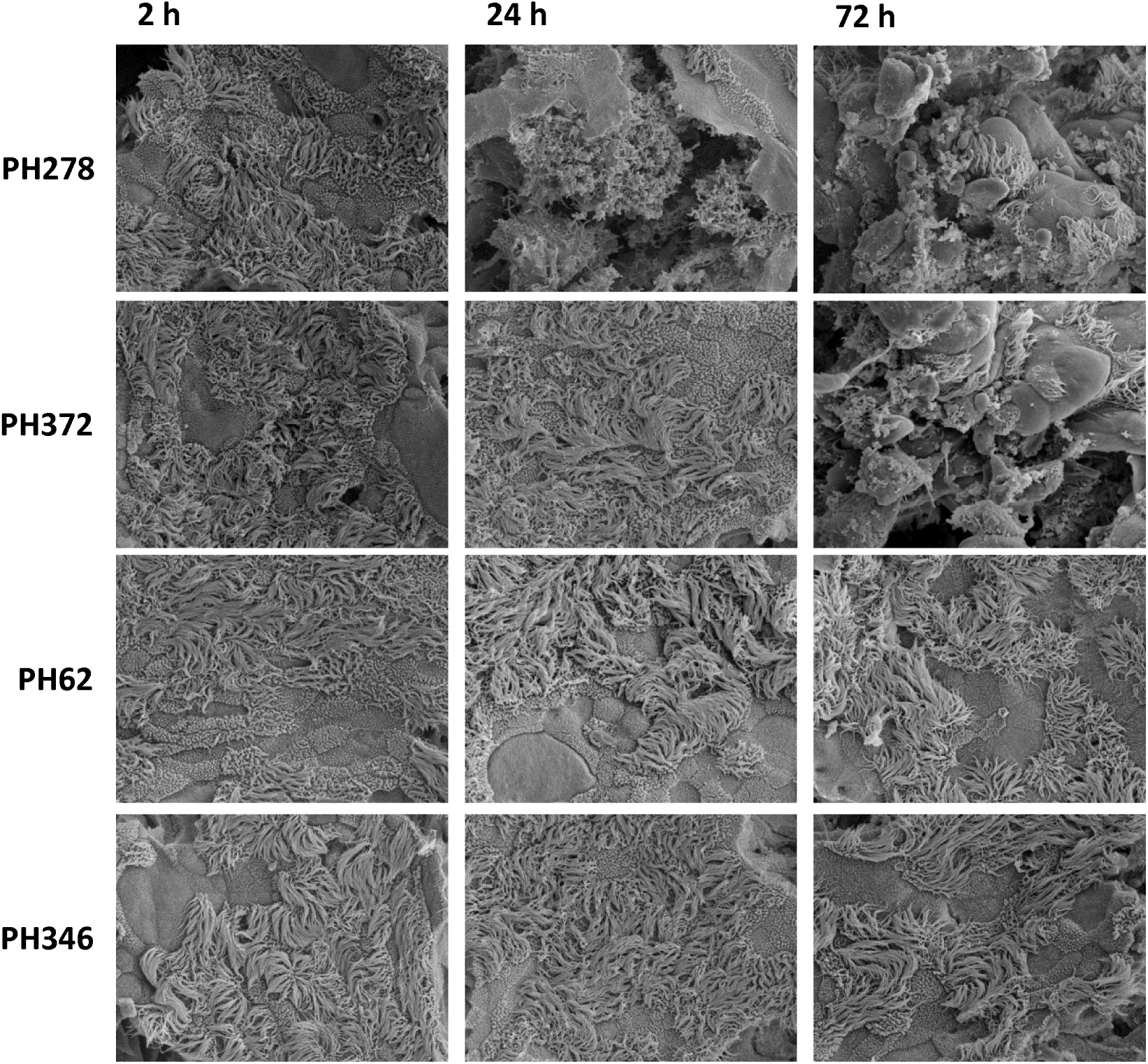
SEM examination of differentiated BBEC culture infected with ovine *M. haemolytica* isolates. BBEC cultures were infected apically with *M. haemolytica* strains PH278, PH372, PH62 or PH346 (2.5 × 10^7^ cfu/insert) at day 21 post-ALI. At 2, 24 or 72 hpi, cultures were apically washed to remove unbound bacteria, fixed and examination by SEM (x2500 magnification).

**Supplementary Figure 12.**
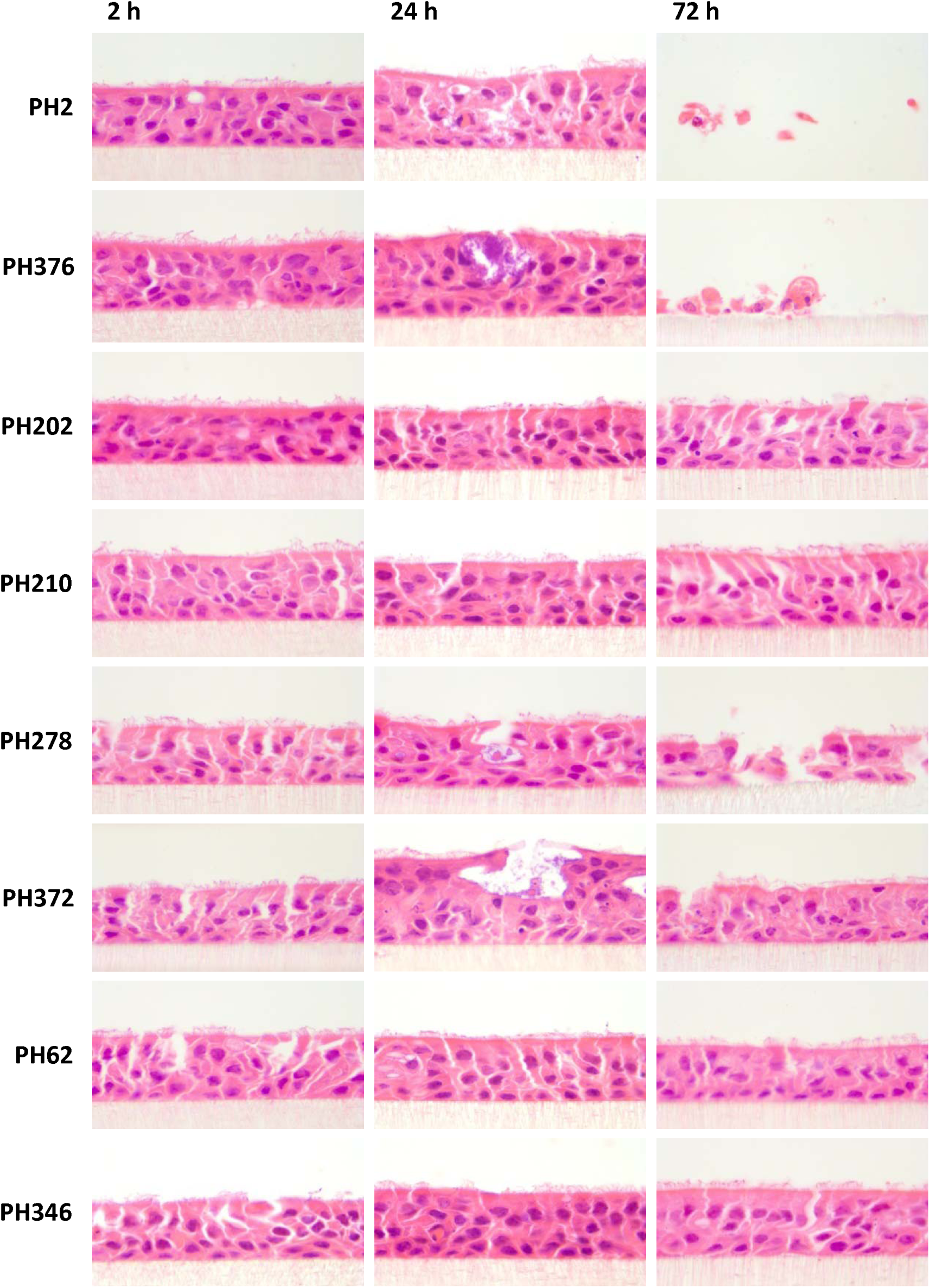
H&E staining of histological sections following *M. haemolytica* infection of differentiated BBEC cultures. BBEC cultures were infected apically with eight strains of *M. haemolytica* (2.5 × 10^7^ cfu/insert) at day 21 post-ALI. At 2, 24 or 72 hpi, cultures were apically washed to remove unbound bacteria, fixed and paraffin-embedded using standard histological techniques. Sections were subsequently cut, deparaffinised and H&E staining was performed (x1000 magnification).

